# Synthetic Phase Variation for Engineered Microbial Consortia

**DOI:** 10.1101/2025.08.25.672192

**Authors:** Matthieu F. Kratz, Richard M. Murray, Michael B. Elowitz

## Abstract

Some biochemical functions can be performed more efficiently when split into different tasks, each performed by distinct strains within a microbial consortium. Due to the distinct growth dynamics of each strain, uncontrolled consortia have unstable composition dynamics, leading to the rapid loss of the community level function. Several approaches have been developed to stabilize consortia, with most relying on communicated-mediated growth and death feedback. These approaches require accurate communication between strains to maintain control, something which is not guaranteed under non-well-mixed conditions. As such, these methods are of limited utility in consortia applications with poorly mixed environments e.g. industrial scale bioreactors or soil. Here, inspired by microbial phase variation dynamics, we introduce an alternative, communication-free approach in which a set of genetically identical cells switch stochastically between distinct phenotypes. In this scheme, the population composition is dynamically stable and determined by the rates of transitions between states. Mathematical modeling indicates that this approach can stabilize consortia. Experimentally, we used reversible DNA inversions catalyzed by serine recombinases to implement a dynamic consortium. We then characterized the dynamic properties of the consortium at the single cell and bulk levels, and demonstrated control in 2- and 3-state consortia. These results provide a composition control approach that does not rely on cell to cell communication, providing a foundation for deployment of engineered consortia in complex, and sometimes non-mixed, environments such as industrial scale bioreactors or the human gut.

## Introduction

Consortia are communities composed of multiple distinct cell types^1–3^ that coordinate their individual behavior to achieve a community-level function. Consortia could implement more complex functions than single strain systems, and enable applications in metabolic engineering and biomanufacturing^4–8^. To this end, a major research goal in microbial synthetic biology has been to extend synthetic gene networks to the consortia level. The standard way to create a consortium is to co-culture several distinct microbial strains implementing separate parts of an overall function^1–3,7^, such as metabolic synthesis of a chemical (Fig. 1a, left). In such systems, optimizing the overall consortium function requires a mechanism to optimize and maintain stable strain composition^1,5,9^. One approach is to use naturally symbiotic strains. However, many applications favor highly engineered and specialized lab strains^10,11^. As a result, even small differences in the growth rates of the individual strains can drastically shift the composition^2,3,7^ (Fig. 1b, left).

**Figure 1.**
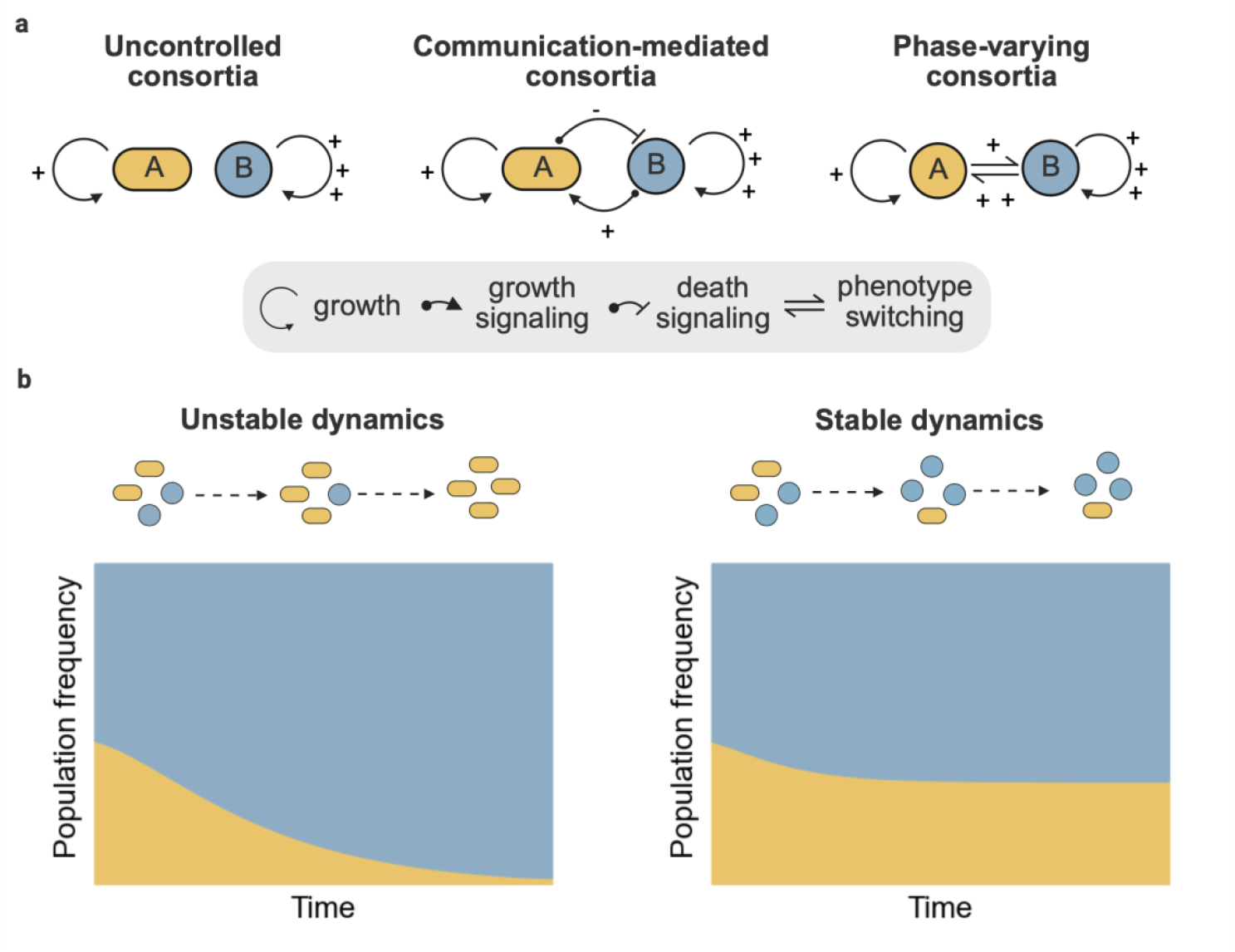
Mechanisms and dynamics of microbial consortia implementations. **a** Schematics of different microbial consortia implementations: (left) no composition control, (middle) communication-based composition control, where growth and death are mediated via cell to cell signalling, and (right) phase-varying consortia, where each cell stochastically switches between phenotypic states. **b** Bulk population dynamics of consortia implementations. In an uncontrolled system, the fastest growing strain dominates the culture (unstable dynamics). Both the communication-mediated and phase-varying consortia balance all strains, achieving stable dynamics. Blue and yellow represent distinct states, while shapes (circle, oval) represent distinct strains.

Several approaches have been introduced to control consortium composition. One approach is to establish feedback on population sizes through intercellular signaling systems (Fig. 1a, center, Fig. 1b right)^12–18^. Broadly speaking, each consortium member secretes a member-specific signal that influences the growth or death rates of other members. For instance, auxotrophic *E. coli* mutants were used to create pairs of strains that complement each other’s growth through cross-feeding of essential metabolites^18^. Metabolites secreted from one auxotrophic strain directly stimulate the growth of a complementary auxotrophic strain, favoring co-existence. In this case, the metabolites act as signals and controllers of growth rates. Recent implementations have expanded the repertoire of metabolites used this way^15^ and exploited quorum sensing molecules and toxin-antitoxin systems for communication and death control^12,13,16^, respectively. These approaches allow for inter-species consortia but require accurate communication between members to properly function^1,4^. As such, their performance can be compromised in non-well-mixed settings e.g. industrial bioreactors^19^ or human gut^20^. Furthermore, they rely on a limited set of orthogonal communication channels, making scaling to many members difficult. Hence, these communication-mediated approaches have substantial limitations in terms of their scalability and suitability for non-well-mixed deployment contexts.

Another recent class of approaches addresses this scaling issue by emulating stem cell differentiation dynamics and removing the need for intercellular communication^21–23^. In these approaches, a consortium starts as a set of undifferentiated progenitor cells and progressively shifts to a community of terminally differentiated cells. Recently, *An et al*. used several orthogonal serine recombinases to mediate differentiation into a series of terminal cell types from a single founder cell type^23^. By manipulating the nucleotide sequence of recombinase attachment sites, the precise ratio of the terminal cell types could be manipulated. This approach is scalable to many phenotypes, but is limited by depletion of the progenitor pool and lacks closed-loop feedback control of overall cell fate proportions.

What approaches could allow for scalable and continual composition control without relying on intercellular communication? In the natural process of phase variation, cells stochastically and reversibly transition between distinct phenotypic states^24,25^, each conferring distinct fitness benefits and costs. Phase variation facilitates survival in environments with abrupt and stochastic changes in selection pressure^24–26^. Here, we implement a synthetic version of phase variation in which a single strain encodes a set of mutually exclusive states, with each individual cell stochastically and reversibly switching between these states (Fig. 1a, right, Fig. 1b right). By controlling state transition rates, the system allows tuning of the equilibrium distribution. In this approach, there is no requirement for intercellular communication and cells have access to all states at all times. These features make this system amenable to continuous control of composition.

We first used mathematical modeling to demonstrate that the dynamics of phase variation can stabilize otherwise unstable consortium configurations. We then designed circuits that exploit reversible recombinase flipping to switch between mutually exclusive phenotypic states. Next, we experimentally constructed this design, showing it can balance the abundances of two distinct states. Finally, through the addition of a secondary orthogonal recombinase, we demonstrated that synthetic phase variation can be scaled to control the proportions of three distinct states. This work establishes a communication-free, scalable architecture for continuous control of consortia composition, opening the door to applications of consortia in otherwise inaccessible contexts.

## Results

### Phase variation stabilizes otherwise unstable consortium *in silico*

Under what circumstances can stochastic switching establish a stable consortium? To address this question, we constructed a simple mathematical model representing the population sizes of cells in two states, denoted *A, B* in a well-mixed, chemostatic environment. Nutrients (*S*) are supplied, and cells and media are removed, at the dilution rate, *D*. Cells divide at rate *µ*. In general, cells compete for the limited nutrient supply.

Using this framework, we first examined the case of an uncontrolled consortium with two competing strains, each fixed in a distinct state, *A* or *B*. For example, if strain *B* grows at a 30% faster rate than strain *A*, and both strains are seeded at equal concentrations, the combination of growth rate differences and limiting nutrient leads strain *B* to progressively overtake the mixed culture (Fig. 2a). Thus, in the absence of additional control mechanisms, the consortium is unstable.

**Figure 2.**
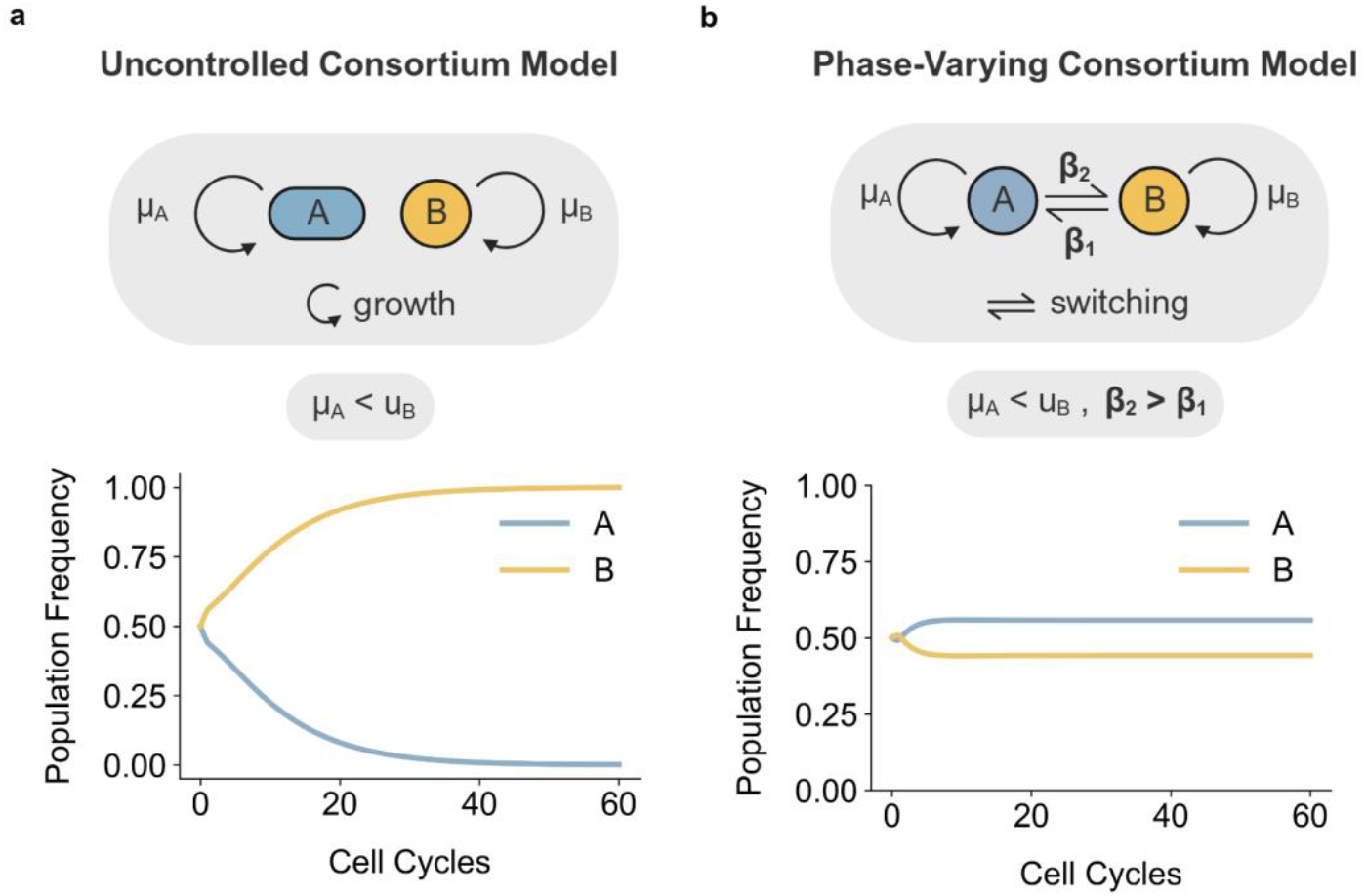
Phase variation stabilizes consortium composition. **a** Uncontrolled consortium model schematic and simulation results. **b** Phase-varying consortium model schematic and simulation results. Presented results and parameters are non-dimensionalized and scaled to the growth rate of strain/phenotype *A*. Further details of models, parameters and simulations can be found in the modeling supplement.

Phase variation mitigates this issue. To explore this, we added a secondary layer of phase variation dynamics to the model. In this case, the two phenotypic states, *A* and *B*, interconvert, with switching rates^27–29^ β_1_ or β_2_ for *B*-to-*A*, or *A*-to-*B* transitions, respectively (Fig. 2b, modeling supplement). Transitions compensate for the growth advantage of *B*, allowing for stable coexistence of both states (Fig. 2b) with a population ratio, *r* (Equation 1), that depends on the switching rates, dilution, and growth rates:

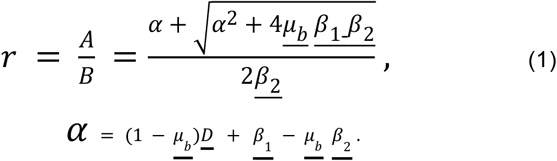

Thus, *r* takes on a unique population ratio that can be modulated by tuning switching rates (Supplementary Fig. 1). Overall, this model demonstrates how phase variation can force co - existence in an otherwise unstable consortia and allow to tune the relative abundance of consortia members.

### A recombinase circuit can drive 2-state phase variation dynamics

Having demonstrated that phase variation dynamics can stabilize consortia *in silico*, we sought to design an experimental circuit that implements phase variation. The ideal circuit would rely on well-characterized components with multiple orthogonal variants to enable scalable engineering of systems with multiple states. We designed a DNA cassette which encodes two mutually exclusive transcriptional states, one corresponding to YFP^30^ expression and the other to mTurquoise^31^ expression (Fig. 3a). Both fluorophores were tagged with the ssrA degradation tag, shortening their half-lives to accelerate circuit readout dynamics^32^. To allow transitions, we also incorporated serine recombinases, viral proteins that catalyze site specific DNA rearrangements^33^. We flanked the cassette with two antiparallel Bxb1 serine recombinase attachment sites. When Bxb1 binds to these sites, it catalyzes inversion of the intervening DNA sequence, switching expression from YFP to mTurquoise. This reaction can be reversed by Bxb1 in complex with its cognate reverse directionality factor (RDF)^34^. By regulating the concentrations of Bxb1 and its RDF, one can influence the frequency and bias of these flipping reactions^35^. Analogous constructs have been previously used as rewritable DNA storage^36–38^ or to tune expression variation of a single gene^39^.

**Figure 3.**
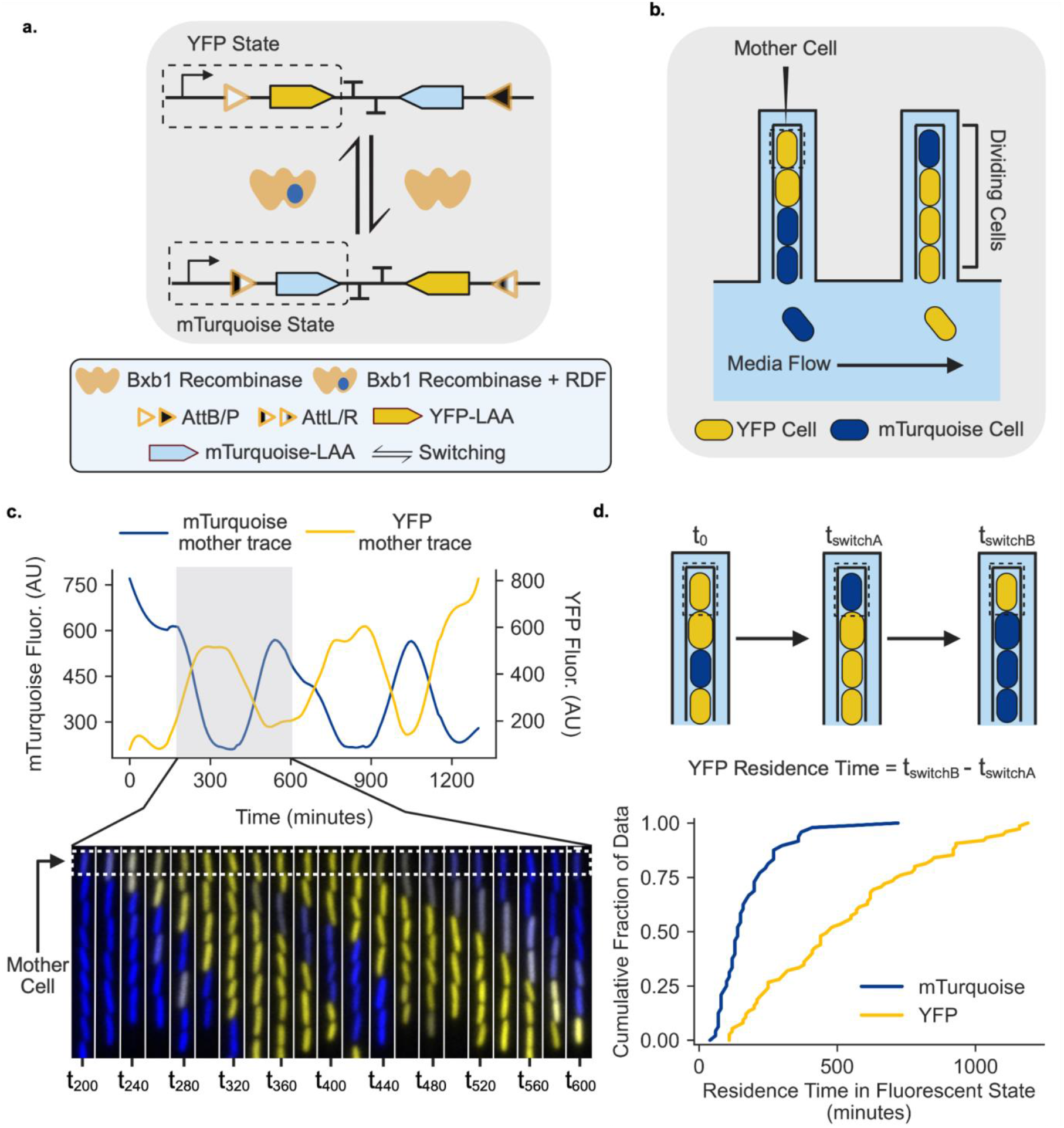
A recombinase circuit successfully recreates single cell phase variation dynamics. **a** Recombinase switching construct schematic. The arrows represent state transitions, with Bxb1 or the Bxb1-RDF complex catalyzing transitions. Bxb1 recombinase was destabilized using the LAA ClpXP degradation tag^32^. This construct was integrated as a single copy in the genome. Bxb1 and RDF were expressed under inducible control from a separate plasmid. **b** Mother machine schematic. The cell at the end of the growth channel (mother cell), remains in the channel indefinitely and can be tracked for many generations **c** Mother machine kymograph and single mother cell fluorescence trace. Successive images in the kymograph are separated by twenty minute intervals. The highlighted grey section in the trace corresponds to the time frame associated with the kymograph movie. **d** Schematic defining residence times and cumulative distribution of mother cell residence times. Data were collected by manual observation of mother cells.

After constructing this circuit, we experimentally probed its single cell dynamics using the ‘mother machine’ microfluidic device^40–44^ (Fig. 3b). This device consists of a series of growth channels perpendicular to a trench through which fresh media flows, providing a chemostatic environment and allowing long term monitoring of individual “mother” cells at the ends of each channel. This device thus allows one to probe the long term switching dynamics of single cells under steady state conditions.

When observed in the mother machine, individual cells stably expressed a single fluorophore for multiple cell cycles, while occasionally switching to the other state. We can see an example of this switching process in Fig. 3c, in which a cell switches from mTurquoise to YFP fluorescence (Supplemental Movie 1). It is important to note that in this instance, the mother cell seems to spend relatively little time in a mixed YFP-mTurquoise state, existing primarily in one of the two states exclusively.

To further probe the switching dynamics, we tracked the distribution of residence times, defined as how long the cell spends in a given state before switching (Fig. 3d). Both states displayed a wide distribution of residence times. Because the switching rates were unequal, cells spent more time in the YFP state (residence time of 521 min., s.d. 300 min) than the mTurquoise state (174 min., s.d. 119 min). Furthermore, consistent with the kymograph observations, the YFP and mTurquoise states were exclusive, with cells spending relatively little time in transitional states expressing both proteins. The residence times were exponentially distributed for both the mTurquoise and YFP states, but with different time constants, consistent with our expectation of a generative Poisson process for switching (Supplementary Fig. 2).

### Varying expression of recombinase elements enables bulk composition control

We next sought to tune the state composition of the population. We placed Bxb1 and Bxb1 RDF under the control of salicylate (sal) and Isopropyl-β-d-thiogalactoside (IPTG) inducible promoters^45^, respectively, on a p15a plasmid (Fig. 4a). With these constructs, the transition rates that govern the bulk cell population dynamics should be tunable, allowing for bulk-scale control of composition.

**Figure 4.**
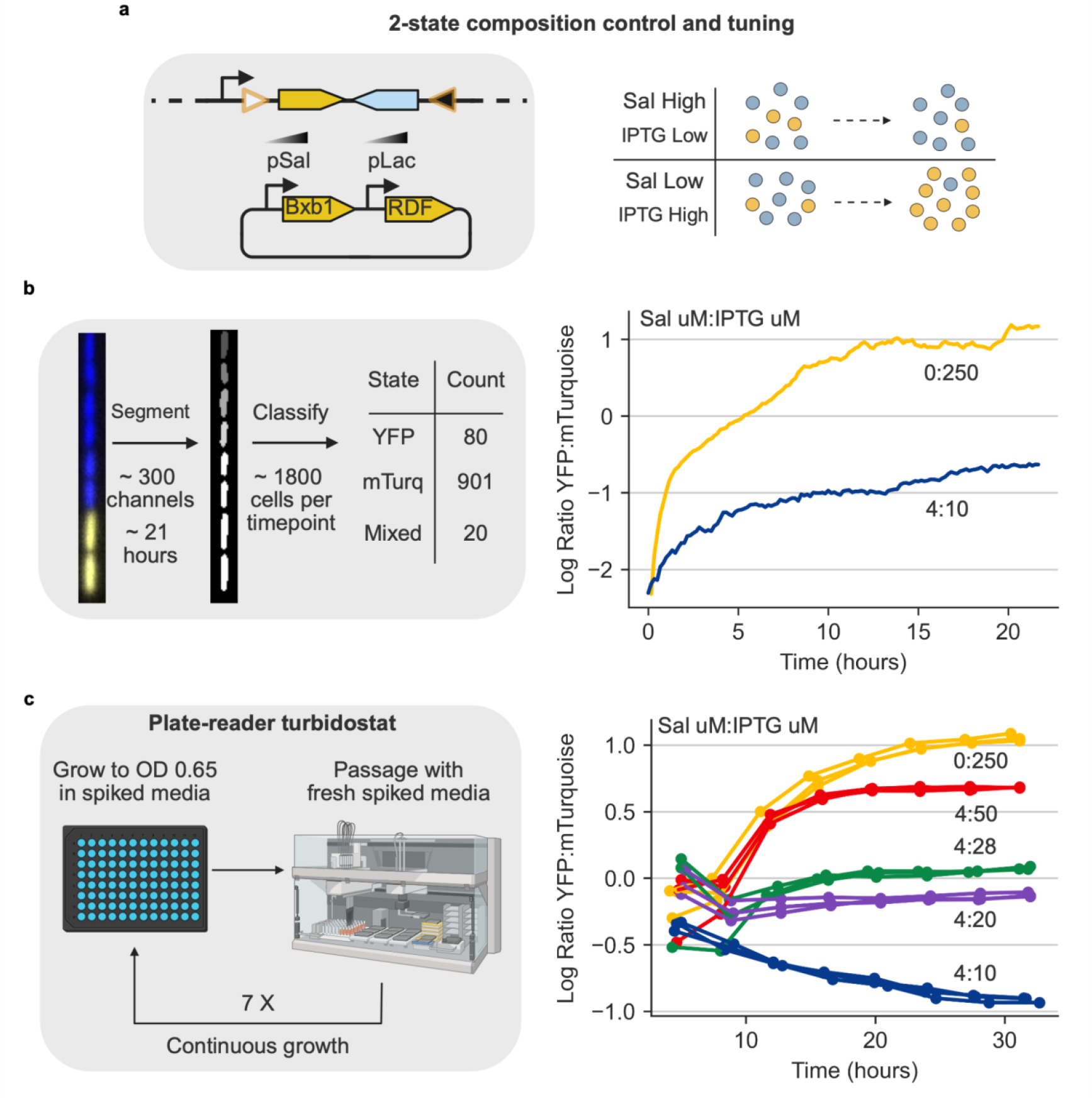
Synthetic phase variation enables bulk composition control in diverse growth contexts. **a** 2-state composition control schematic. The recombinase switching construct was located in a single copy on the genome and the recombinase elements were expressed from a p15a plasmid. Bxb1 is controlled by a salicylate inducible promoter (pSal) and its cognate RDF is controlled by an IPTG inducible promoter pTac. By varying inducer levels, the steady state composition of the system should be tunable. **b** Mother machine processing pipeline and bulk population dynamics. Cells are sorted into YFP and mTurquoise categories based on the median fluorescent signal ratio of the relevant channels. Traces start in a heavily mTurquoise biased state due to leak associated with the Bxb1 recombinase. The inducer concentrations used for each experiment are labeled next to each trace. Individual population traces can be found in Supplementary Fig. 2-3. **c** Plate reader turbidostat setup and bulk population dynamics. Traces of the same color are cultured in the same media and expected to end at the same final composition. The inducer concentrations used for each experiment are labeled next to each trace. Due to the dynamics of degradation-tagged fluorophores in batch cultures, the maximum value for each fluorophore channel in each cycle was used to calculate composition. Signals are normalized to the max fluorescence possible for a given channel e.g. YFP is normalized to an all YFP culture.

We used the mother machine to observe population dynamics under YFP-biased (0 µM sal, 250 µM IPTG) and Turquoise-biased (4 µM sal, 10 µM IPTG) inducer conditions. The circuit was strongly biased to the mTurquoise state in the absence of inducers due to basal Bxb1 expression. Upon induction, the distribution of cell states rapidly shifted to a new steady-state composition (Fig. 4b, Supplementary Fig. 3-4). Within ∼19 hours both conditions reached 95% of the steady-state composition value. Additionally, we tracked the frequency of mixed-state cells during these runs (Supplementary Fig. 5) and noted that at steady state, the frequency of these cells remained ∼5%, further supporting our previous findings that most cells exist in a single expression state. Together, these results indicate that Bxb1 and RDF expression levels can tune the population composition.

To rapidly explore a broader range of inducer concentrations, we switched to a turbidostat-plate reader setup that allows greater multiplexing (Fig. 4c). The non-chemostatic plate reader conditions also provided an opportunity to examine circuit operation in a distinct, batch culture growth context.

For plate reader experiments, we explored a matrix of 3 initial compositions, each induced with one of 5 different inducer concentrations, for a total of 15 individual culture trajectories (Fig. 4c, Supplementary Fig. 7-11). For each inducer concentration, cultures approached the same final steady state distribution regardless of initial conditions, and did so over a similar timescale (Fig. 4c). Final compositions spanned a ∼100-fold range of population ratios, and were tunable within this range. In a negative control experiment, we performed a similar analysis on a 2-strain (non-switching) system. In this case, cultures were overtaken by the YFP expressing strain (Supplementary Fig. 6), indicating that the circuit is required to stabilize mixed compositions.

The time to reach 95% of the steady state was noticeably longer in the plate reader (∼19 h) compared to the mother machine (∼26 h). However, the final compositions were similar between the two environments — 1.08 vs. 1.05 for the YFP-biased condition and -0.91 vs. -0.66 for the mTurquoise-biased condition. Taken together, these results demonstrated continuous control of bulk composition dynamics.

### A second recombinase enables 3-state phase variation

Many consortium applications primarily use two strains, but recent use cases in metabolic engineering employed as many as four distinct strains^46^. This work provokes the question of whether synthetic phase variation can scale to larger numbers of states^47,48^. In fact, incorporation of a second orthogonal recombinase (blue) can enable 3-state switching. As shown in Fig. 5a, we added a third transcriptional state downstream of the 2-state unit, and flanked the construct with the attachment sites of the second recombinase. The resulting system is predicted to generate four genetic states, one for state A, one for state B and two for state C (C_A_ and C_B_). These two C states both express the C gene and are therefore phenotypically equivalent. However, they respond differently to the second (blue) recombinase, with C_A_ and C_B_ respectively transitioning to states A and B. In this design, the first recombinase activity (yellow) determines the ratio of A to B population sizes, while the second recombinase (blue) determines the (A+B) to C ratio. We refer to the system as the 3-state circuit based on the number of distinct phenotypic states.

**Figure 5.**
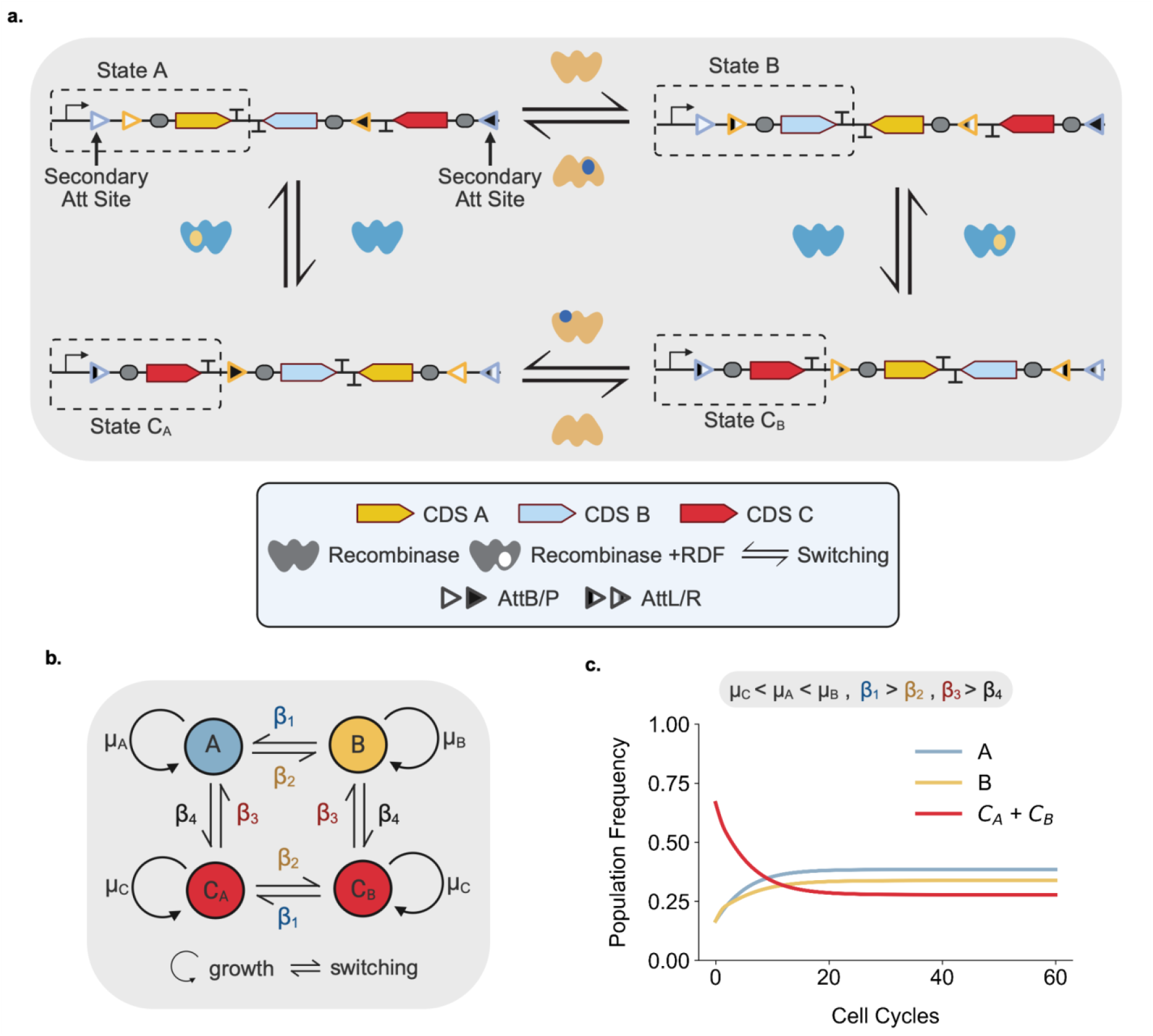
A second recombinase allows 3-state switching. **a**, Recombinase elements (include Att sites) are colored according to their cognate recombinase. Arrows represent transitions between states, with the relevant recombinase complex catalyzing these transitions next to each arrow. The active transcriptional state of the system is surrounded by a dotted box. Note C_A_ and C_B_ are two separate DNA states but transcriptionally are the same state. **b** 3-state phase varying consortium schematic. Note that certain transition rates catalyze multiple distinct transitions e.g. β_1_ switches B to A and C_B_ to C_A_. **c** 3-state model simulation dynamics. Presented results and parameters are non-dimensionalized and scaled to the growth rate of strain/phenotype C. Further details on modeling and simulations can be found in the modeling supplement.

To understand composition control properties of the 3-state circuit, we simulated its dynamics using the framework in Fig. 2 (Fig. 5b, modeling supplement). State-switching in the 3-state architecture stabilizes configurations that would otherwise be unstable (Fig. 5c). Depending on switching rates, the circuit can access the full repertoire of possible cell state compositions (Supplementary Fig. 12). Overall, these mathematical modeling results imply that our design can be scaled beyond the control of 2-states.

### A recombinase circuit experimentally demonstrates 3-state phase variation

To implement the 3-state circuits, we added a second orthogonal recombinase and RDF, TP901^49^ together with a third fluorescent protein, mScarlet3^50^. TP901 and its cognate RDF were placed under the control of the lux and cin inducible promoters^45^, respectively and assembled alongside the Bxb1 elements (Fig. 6a). We examined the system in the mother machine, using similar growth conditions to the 2-state system (Fig. 6b, Methods). Cells switched across all three fluorophore states, presenting stable expression of each fluorescent state. However, in contrast to the 2-state system, the 3-state circuit imposed a larger burden on cell physiology, leading to filementation and cell death (Supplemental Movie 2). This was partially remedied by reducing the copy number of the recombinase elements by moving the circuit from a p15a plasmid to a genomic location (pOSIP T-site)^51^. This alteration successfully reduced burden. It also markedly increased the single cell switching timescale, consistent with lower recombinase expression levels (Supplemental Movie 3-7). Nonetheless, the 3-state system exhibited active switching among three stable, heritable expression states.

**Figure 6.**
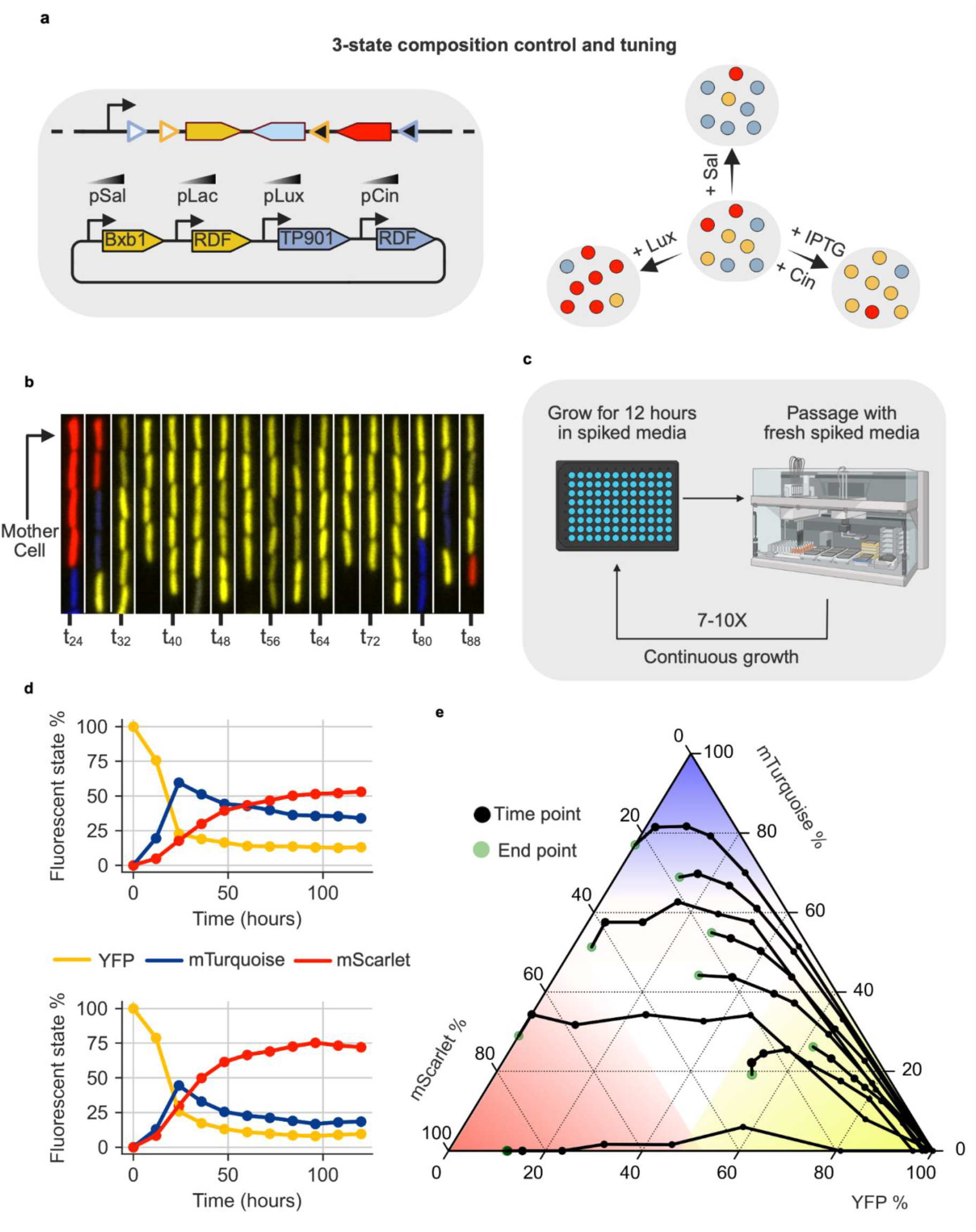
The 3-state design retains the single cell and bulk dynamics of the 2-state system. **a** 3-state composition control schematic. Recombinase elements (include Att sites) are colored according to their cognate recombinase. The 3-state switching construct is integrated as a single copy on the genome. Bxb1 is controlled by a salicylate inducible promoter (pSal) and its cognate RDF is controlled by an IPTG inducible promoter pTac. TP901 is controlled by a Lux AHL (pLux) inducible promoter and its cognate RDF is controlled by a Cin AHL inducible promoter (pCin). By varying inducer levels, the steady state composition of the system should be tunable. The copy number of the recombinase element varies based on the experiment. **b** Mother machine kymograph. Successive images are separated by forty minute intervals. **c** Liquid handler passaging setup. **d** Select stable 3-state time course traces. Fluorescence values have been normalized to the max possible fluorescence signal for the given channel. **e** 3-state space exploration traces. Fluorescence values have been normalized to the max possible fluorescence signal for the given channel. As typical for a ternary diagram, the ratio of all three channels is used to calculate positions of individual trajectories. Each point in a given trajectory is separated by a fixed growth cycle of 12 hours.

Finally, we characterized composition control at the bulk level. We initiated cultures from an all-YFP initial composition. We identified several inducer conditions that enabled stable, long-term coexistence (∼120 hours) of all states (Fig. 6d, Supplementary Fig. 23-24). This contrasts with a non-switching control system, where the mScarlet strain rapidly takes over the mixed culture (Supplementary Fig. 25). Interestingly, the two switching processes in our system operate at different timescales, with the YFP to mTurquoise (Bxb1) process reaching equilibrium noticeably faster than the (YFP + mTurquoise) to mScarlet process. These experiments demonstrated that synthetic phase variation circuits can create a stable set of compositions in an otherwise unstable 3-state system.

Finally, having investigated stable compositions, we asked what transient composition trajectories were accessible. We used a similar passaging setup, but assayed a wider variety of inducer concentrations over fewer passages (Supplementary Table 1). Under these conditions, increasing salicylate concentration (Bxb1) increased mTurquoise signal and increasing lux AHL (TP901) increased mScarlet signal, indicating that the system continued to function as expected. The 3 - state system traversed a broad variety of compositions (Fig. 6e, Supplementary Fig. 13-22). Cultures transiently populated compositions heavily biased towards individual states (closer to vertices), as well as mixed compositions (closer to grey center triangle). Depending on the conditions, cells traversed the composition space at different velocities. These experiments demonstrate that we can transiently pass through a wide variety of compositions.

## Discussion

The ability to establish defined distributions of cell states within a population would facilitate the creation of stable microbial communities. It has, however, remained unclear if a recombinase approach could tune phenotypic variation for multiple distinct states, and improve upon the tunability and scalability of toggle switches^48,52,53^ and other previous approaches^14,22,23^. Here, we demonstrated granular, straightforward control of two to three distinct phenotypic states. Further, this strategy could be scaled to higher numbers of states by adding additional recombinases. Thus, it should be possible to engineer multi-state consortia within a single strain.

There are several limitations of this work. First, recombinases place a burden on cell physiology^54–56^. Better understanding this burden and developing ways to circumvent it will be important for its deployment in applications. Specifically, characterizing the interplay between recombinase expression, switching rates, bulk dynamics and burden will be critical. Optimizing burden is a particular concern as a given composition control method should be as low-burden as possible to minimize interference with consortia function. However, the circuits introduced here displayed high levels of burden in the mother machine, with noticeable cell death and filamentation, especially in the 3-state system (Supplemental Movie 2). These effects were reduced but not eliminated by reducing the copy number of the recombinase genes. In this architecture, the number of unique controllable states scales linearly with the number of distinct recombinases. There are over 60 characterized serine recombinases^35^, many of which are orthogonal but the larger issue relates to the burden associated with each additional recombinase. This could be partially addressed through mutant recombinase attachment sites that switch at faster rates than their wild-type counterparts^57^, reducing the required recombinase expression level, while maintaining higher switching rates. Another fundamental limitation of our approach is shared with the terminal differentiation approach: the cell must be capable of expressing each phenotype in the system. This prevents use of multi-species consortia and runs counter to the typical strain specialization workflows of metabolic engineering^8,11^.

Synthetic phase variation could be expanded in several ways. First, one could decouple the core state switching circuit from the target genes it controls. Specifically, fluorescent protein genes could be replaced with transcriptional activators that would in turn regulate expression of functional transcriptional units on separate genetic constructs. Those transcriptional units would not need to be expressed as single copies. This modification would improve modularity, allowing for easy swapping of functional target genes. Second, a unique element of our circuit is that it could be implemented in mammalian systems, where recombinase activity has previously been demonstrated^58^. Further, unlike communication-mediated composition control, the circuit introduced here does not rely on synthesis or active import and export of metabolites, making it theoretically more host-agnostic than other approaches. It will be interesting to find out whether the circuit can be modified to operate in human cells for research and applications. The approach introduced here could potentially be implemented in non-model microorganisms for rapid development of non-model organism consortia, e.g. *Pseudomonas* species for soil^59^ and *Bacteroides* for gut^60^.

Synthetic phase variation circuits enable several specific applications. Because cells retain access to all states and control can be maintained without intercellular communication, composition control should be spatially invariant and deployable in non-well-mixed, even static, environments. For instance, bioprocessing has benefited from use of consortia^46,61–63^. However, at the industrial scale many fermentations have issues with spatial heterogeneity and mixing^19^. This interferes with the use of communication-mediated consortia; our approach would be able to maintain control in this context. Similarly, gut microbiome engineering could be heavily enabled through the use of consortia, with different phenotypes interacting with specific niches to cause persistent, large scale shifts in the microbiome^64^. However, the gut displays erratic and irregular mixing, favouring the use of our approach to implement consortia in the gut microbiome ^20^. Another suitable use of our system may be in the context of biomaterials. Specifically, several biomaterials systems use varying ratios of different strains/phenotypes to generate materials with differing mechanical properties^65,66^. In our framework, a single cell would be able to generate all phenotypes, implying one could regenerate a biomaterial from a single cell sample. This regeneration phenomena would be infeasible to recreate using the communication-mediated or terminal differentiation approaches.

Stochasticity is often perceived as a nuisance that degrades the performance of synthetic biological circuits. In this case, stochasticity is essential to generate stable, accurate composition at the population level. This design provides an example of how naturally inspired circuits can be used to exploit unique biological phenomena to generate novel engineered functions.

### Strain and plasmid construction

A list of all fragments, constructs and strains used in this study can be found in Supplementary File 1. The associated sequences have been uploaded as part of the supplemental files. The MG1655 Marionette strain was used as the base strain for all experiments in this study. Genomic integrations for this study were performed using the pOSIP clonetegration system. For mother machine experiments, knockouts of the MotA gene were necessary and were performed using Lambda Red Recombineering.

All plasmids were assembled using 3G assembly^1^, sourcing parts from Murray Biocircuits Part Library, an extended version of the CIDAR MoClo extension part kit. To perform pOSIP clonetegration of switching constructs, an approach alternating use of bsaI (NEBuilder® HiFi DNA Assembly Master Mix) and bsmbI (NEBridge® Golden Gate Assembly Kit (BsmBI-v2)) to catalyze golden gate assembly constructs was employed. Other pOSIP clonetegrations were performed using 3G assembly.

### Media and Reagents

For plate reader experiments, cells were grown in M9CA 1% glucose (Teknova) with 25 μg/ml chloramphenicol and 50 μg/ml kanamycin. For mother machine experiments cells were grown in M9CA 0.5% glycerol with 25 μg/ml chloramphenicol, 50 μg/ml kanamycin and 50 μg/ml hygromycin. Unlike the glucose media, the glycerol media was not purchased from a vendor and made in the wetlab. 1L of this media was produced using 200 ml of 5X M9 Salts, 10 ml 50% glycerol solution (Teknova), 10 ml 100X casamino acid solution, 1 ml 1000X thiamine-HCL solution, 200 µl 5000X MgSO_4_ solution and 100 µl 10000X CaCl_2_ solution. The quantities of each chemical and solvent required to make these stock solutions can be found in the table below.

| Solution name | Chemical name and quantity | Solvent name and quantity |
| --- | --- | --- |
| 5X M9 Salts | 56.4 g M9 Salts | 1L distilled water |
| 100X Casamino acid | 20 g Casamino acids | 200 mL distilled water |
| 1000X Thiamine | 10 mg Thiamine-HCl | 20 mL distilled water |
| 5000X MgSO<sub>4</sub> | 2.465 mg MgSO<sub>4</sub>·7H<sub>2</sub>O | 10 mL distilled water |
| 10000X CaCl<sub>2</sub> | 1.10 mg CaCl<sub>2</sub>·2H<sub>2</sub>O | 10 mL distilled water |

### Mother machine experiment preparation

Microfluidic ‘mother machine’ devices were manufactured in the Jun Lab at UCSD according to the protocol from Wang et al.^2^ Prior to loading cells, the device was annealed to a WillCo-dish (Glass Bottom Dish, HBST-5040) via high power plasma cleaning. Once annealed, the device was loaded with 10 µl of 0.5 mg mL^-1^ BSA (Bovine serum albumin).

From an overnight culture of the relevant strain grown in M9CA Glycerol (w/ antibiotics), 20 µl of culture was diluted into 1 ml of fresh M9CA Glycerol (w/ antibiotics) and grown at 37 °C for 4 hours. After 4 hours, the culture was centrifuged and resuspended to concentrate cells 10-fold. 10 µl of this concentrated culture was then loaded into the BSA-treated mother machine device main trench. To load cells into the channels of the mother machine, the device was then placed (via custom adapters) into a centrifuge and spun for two minutes on each side of the device, loading cells into channels on both sides of the main trench. Once cells were loaded, small volumes of inducer spiked media were loaded from the syringe into the mother machine device to clear the main trench of cell debris and provide media for cells.

### Mother machine analysis

To correct temperature related drift during the mother machine experiments, the ‘Correct 3D Drift’ plugin in FIJI^3^ was employed for each FOV. Stabilized FOVs were then cropped and resized to ensure identical pixel dimensions across FOVs. FOVs were then analyzed and segmented in the napari-mm3 software^4^ developed by the Jun Lab.

The subsequent described steps were performed in python and the scripts applying these steps to the segmented dataset can be found at the following GitHub repository. With the segmented outputs of the napari-mm3 pipeline, the cells masks were applied to the YFP and mTurquoise fluorescent channels. After background subtraction and normalization to the maximum signal of each fluorophore, intensities for each cell were summed. If this sum was below 0.25, the cell was assumed to be non-fluorescent and not included in further analysis. Cells that passed this screening step had the ratio of normalized mTurquoise intensity to normalized YFP intensity calculated. Using thresholds based on observations of movies, a cell was considered mTurquoise if this ratio was above 2, YFP if this ratio was below 0.5 and mixed if between these two values. To calculate the steady-state composition, we used the average log_10_(YFP/mTurquoise) signal for the last 3 hours of each experimental sample. To calculate the time to 95% steady-state, we located the first time point where log_10_(YFP/mTurquoise) was within 5% of the aforementioned steady state.

For the mother cell trace in Fig 3., on the stabilized movie of the relevant trench, a region of interest in FIJI was drawn that encapsulated exclusively the mother cell. The ‘Measure Stack’ function was then used to collect intensities in this ROI. The max values of each channel across time were then smoothed using the Savitzky-Golay filter implementation in SciPy^5^.

### Composition control plate reader experiments (2-state)

Unless otherwise specified, all cultures were grown in 96 square well plates (96 well MatriPlate™ clear bottom), in M9CA Glucose with kanamycin and chloramphenicol, at 37 °C using a plate reader (Biotek H1M). From an overnight culture of the 2-state system, three separate wells with different spiked media were seeded and grown overnight. These three wells were then spun down to remove inducers and resuspended in fresh media. 65 ul of these resuspended cultures were then added to 299 ul of fresh unspiked media and grown in a plate reader until they reached an OD_700_ of 0.65. At this OD, the liquid handler (Hamilton STARlet Liquid Handling Robot) removed the culture plate and seeded a new culture for each of the three wells. These new cultures were composed of 299 µl spiked media and 64 µl of the previous culture. The culture plate was then replaced into the plate reader, repeating this culture seeding process once the newest culture reached an OD_700_ of 0.65.

Due to the time-varying, pulsatile production of the deg-tagged fluorophores in batch culture, the max YFP and mTurquoise fluorescent signal pulse in a given growth well was taken as the abundance of a given state. After background subtraction, these values were used to calculate the ratio of YFP to mTurquoise signal. To calculate the steady-state composition, we used the average log_10_(YFP/mTurquoise) signal for the last timepoint across the three replicates for a given induction condition. To calculate the time to 95% steady-state, we located the first time point where log_10_(YFP/mTurquoise) was within 5% of the aforementioned steady state.

### Composition control plate reader experiments (3-state)

From an overnight culture of the 3-state system, a single well with spiked media was seeded and grown in the plate reader at 37°C. After 12 hours, the plate was removed and 2 µl of cells from the grown well was transferred to a new well with 200 µl fresh spiked media, maintaining the same concentrations of spiked inducers in these new wells. This lower dilution ratio was selected as it seemed to cause a marked improvement in cell health and circuit function. For stable, long time courses (Fig 6d), this dilution occurred 9 times for a total observed growth time of approximately 10X12 hrs. For space exploration time courses (Fig 6e), this dilution occurred 7 times for a total observed growth time of approximately 7X12 hrs.

Due to the time-varying production of the deg-tagged fluorophores in batch culture, the max YFP, mScarlet and mTurquoise fluorescent signal in a given growth well was taken as the abundance of a given state. Due to drastically differing max intensities of each state, after background subtraction, the max signal was normalized to the maximum possible intensity of the relevant fluorescent state. These max intensities were determined by completely biasing the system to the relevant fluorescent state. These subtracted and normalized values were then used to plot the ternary diagram in Figure 6.

### Data and code for computational analysis

Qualitative descriptions of computational analysis performed are present in each subsection of the methods. Data used for analysis and scripts executing said analysis can be found in the [Github repository](https://github.com/mkratz/Phase-variation) associated with this manuscript.

## Supporting information

Main supplement (figures, tables and modeling)

List of strains, plasmids and constructs

Annotated sequences of plasmids and constructs

## Acknowledgments

We thank John Marken, Manisha Kapasiawala, Ethan Richman and Evan Mun as well as other members of the Murray and Elowitz Lab for their feedback on this manuscript. We also thank the various members of the Jun Lab, especially Michael Sandler and Haochen Fu for hosting Matthieu in their lab for the mother machine experiments. This material is based upon work funded by the Caltech Rosen Bioengineering Center and the Caltech Center for Environmental Microbial Interaction. MBE is an investigator of the Howard Hughes Medical Institute.

## Contributions

MFK conceived and designed the project, performed the experiments, analyzed the data, and wrote the paper, all under the supervision of MBE and RMM.

## References

1. McCarty, N. S. & Ledesma-Amaro, R. Synthetic Biology Tools to Engineer Microbial Communities for Biotechnology. Trends Biotechnol. 37, 181–197 (2019).

2. Brenner, K., You, L. & Arnold, F. H. Engineering microbial consortia: a new frontier in synthetic biology. Trends Biotechnol. 26, 483–489 (2008).

3. Duncker, K. E., Holmes, Z. A. & You, L. Engineered microbial consortia: strategies and applications. Microb. Cell Factories 20, 211 (2021).

4. Roell, G. W. et al. Engineering microbial consortia by division of labor. Microb. Cell Factories 18, 35 (2019).

5. Mittermeier, F. et al. Artificial microbial consortia for bioproduction processes. Eng. Life Sci. 23, e2100152 (2023).

6. Jawed, K., Yazdani, S. S. & Koffas, M. AG. Advances in the development and application of microbial consortia for metabolic engineering. Metab. Eng. Commun. 9, e00095 (2019).

7. Bhatia, S. K. et al. Biotechnological potential of microbial consortia and future perspectives. Crit. Rev. Biotechnol. 38, 1209–1229 (2018).

8. Xin, Y. & Qiao, M. Towards microbial consortia in fermented foods for metabolic engineering and synthetic biology. Food Res. Int. 201, 115677 (2025).

9. Jiang, Y., Wu, R., Zhang, W., Xin, F. & Jiang, M. Construction of stable microbial consortia for effective biochemical synthesis. Trends Biotechnol. 41, 1430–1441 (2023).

10. Sagt, C. M. J. Systems metabolic engineering in an industrial setting. Appl. Microbiol. Biotechnol. 97, 2319–2326 (2013).

11. Sun, J. & Alper, H. S. Metabolic engineering of strains: from industrial-scale to lab-scale chemical production. J. Ind. Microbiol. Biotechnol. 42, 423–436 (2015).

12. Balagaddé, F. K. et al. A synthetic Escherichia coli predator-prey ecosystem. Mol. Syst. Biol. 4, 187 (2008).

13. McCardell, R. D., Pandey, A. & Murray, R. M. Control of density and composition in an engineered two-member bacterial community. 632174 Preprint at 10.1101/632174 (2019).

14. You, L., Cox, R. S., Weiss, R. & Arnold, F. H. Programmed population control by cell– cell communication and regulated killing. Nature 428, 868–871 (2004).

15. Kong, W., Meldgin, D. R., Collins, J. J. & Lu, T. Designing microbial consortia with defined social interactions. Nat. Chem. Biol. 14, 821–829 (2018).

16. Scott, S. R. et al. A stabilized microbial ecosystem of self-limiting bacteria using synthetic quorum-regulated lysis. Nat. Microbiol. 2, 1–9 (2017).

17. Xu, P. Dynamics of microbial competition, commensalism, and cooperation and its implications for coculture and microbiome engineering. Biotechnol. Bioeng. 118, 199–209 (2021).

18. Wintermute, E. H. & Silver, P. A. Emergent cooperation in microbial metabolism. Mol. Syst. Biol. 6, 407 (2010).

19. Lara, A. R., Galindo, E., Ramírez, O. T. & Palomares, L. A. Living with heterogeneities in bioreactors: understanding the effects of environmental gradients on cells. Mol. Biotechnol. 34, 355–381 (2006).

20. Labavić, D., Loverdo, C. & Bitbol, A.-F. Hydrodynamic flow and concentration gradients in the gut enhance neutral bacterial diversity. Proc. Natl. Acad. Sci. 119, e2108671119 (2022).

21. Williams, R. L. & Murray, R. M. Integrase-mediated differentiation circuits improve evolutionary stability of burdensome and toxic functions in E. coli. Nat. Commun. 13, 6822 (2022).

22. Aditya, C., Bertaux, F., Batt, G. & Ruess, J. A light tunable differentiation system for the creation and control of consortia in yeast. Nat. Commun. 12, 5829 (2021).

23. An, B. et al. Synthetic ratio computation for programming population composition and multicellular morphology. 2024.11.26.624747 Preprint at 10.1101/2024.11.26.624747 (2024).

24. Thattai, M. & van Oudenaarden, A. Stochastic Gene Expression in Fluctuating Environments. Genetics 167, 523–530 (2004).

25. van der Woude, M. W. & Bäumler, A. J. Phase and Antigenic Variation in Bacteria. Clin. Microbiol. Rev. 17, 581–611 (2004).

26. Morawska, L. P., Hernandez-Valdes, J. A. & Kuipers, O. P. Diversity of bet-hedging strategies in microbial communities—Recent cases and insights. Wires Mech. Dis. 14, e1544 (2022).

27. Wolf, D. M., Vazirani, V. V. & Arkin, A. P. Diversity in times of adversity: probabilistic strategies in microbial survival games. J. Theor. Biol. 234, 227–253 (2005).

28. Visco, P., Allen, R. J., Majumdar, S. N. & Evans, M. R. Switching and Growth for Microbial Populations in Catastrophic Responsive Environments. Biophys. J. 98, 1099–1108 (2010).

29. Jia, C., Qian, M.-P., Kang, Y. & Jiang, D.-Q. Modeling stochastic phenotype switching and bet-hedging in bacteria: stochastic nonlinear dynamics and critical state identification. Preprint at 10.48550/arXiv.1311.2216 (2015).

30. Pédelacq, J.-D., Cabantous, S., Tran, T., Terwilliger, T. C. & Waldo, G. S. Engineering and characterization of a superfolder green fluorescent protein. Nat. Biotechnol. 24, 79–88 (2006).

31. Goedhart, J. et al. Structure-guided evolution of cyan fluorescent proteins towards a quantum yield of 93%. Nat. Commun. 3, 751 (2012).

32. Jadhav, P., Chen, Y., Butzin, N., Buceta, J. & Urchueguía, A. Bacterial degrons in synthetic circuits. Open Biol. 12, 220180 (2022).

33. Stark, W. M. Making serine integrases work for us. Curr. Opin. Microbiol. 38, 130–136 (2017).

34. Olorunniji, F. J. et al. Gated rotation mechanism of site-specific recombination by ϕC31 integrase. Proc. Natl. Acad. Sci. 109, 19661–19666 (2012).

35. Durrant, M. G. et al. Systematic discovery of recombinases for efficient integration of large DNA sequences into the human genome. Nat. Biotechnol. 41, 488–499 (2023).

36. Yang, L. et al. Permanent genetic memory with >1-byte capacity. Nat. Methods 11, 1261–1266 (2014).

37. Bonnet, J., Subsoontorn, P. & Endy, D. Rewritable digital data storage in live cells via engineered control of recombination directionality. Proc. Natl. Acad. Sci. 109, 8884–8889 (2012).

38. Huang, B. D., Kim, D., Yu, Y. & Wilson, C. J. Engineering intelligent chassis cells via recombinase-based MEMORY circuits. Nat. Commun. 15, 2418 (2024).

39. Hung, M. et al. Modulating the frequency and bias of stochastic switching to control phenotypic variation. Nat. Commun. 5, 4574 (2014).

40. Wang, P. et al. Robust growth of Escherichia coli. Curr. Biol. CB 20, 1099–1103 (2010).

41. Bakshi, S. et al. Tracking bacterial lineages in complex and dynamic environments with applications for growth control and persistence. Nat. Microbiol. 6, 783–791 (2021).

42. Luro, S., Potvin-Trottier, L., Okumus, B. & Paulsson, J. Isolating live cells after high-throughput, long-term, time-lapse microscopy. Nat. Methods 17, 93–100 (2020).

43. Sauls, J. T. et al. Control of Bacillus subtilis Replication Initiation during Physiological Transitions and Perturbations. mBio 10, 10.1128/mbio.02205-19 (2019).

44. Si, F. et al. Mechanistic Origin of Cell-Size Control and Homeostasis in Bacteria. Curr. Biol. 29, 1760-1770.e7 (2019).

45. Meyer, A. J., Segall-Shapiro, T. H., Glassey, E., Zhang, J. & Voigt, C. A. Escherichia coli “Marionette” strains with 12 highly optimized small-molecule sensors. Nat. Chem. Biol. 15, 196–204 (2019).

46. Jones, J. A. et al. Complete Biosynthesis of Anthocyanins Using E. coli Polycultures. mBio 8, e00621–17.

47. Zhu, R., del Rio-Salgado, J. M., Garcia-Ojalvo, J. & Elowitz, M. B. Synthetic multistability in mammalian cells. Science 375, eabg9765 (2022).

48. Wu, F., Su, R.-Q.Lai, Y.-C. & Wang, X. Engineering of a synthetic quadrastable gene network to approach Waddington landscape and cell fate determination. eLife 6, e23702 (2017).

49. Christiansen, B., Brøndsted, L., Vogensen, F. K. & Hammer, K. A resolvase-like protein is required for the site-specific integration of the temperate lactococcal bacteriophage TP901-1. J. Bacteriol. 178, 5164–5173 (1996).

50. Gadella, T. W. J. et al. mScarlet3: a brilliant and fast-maturing red fluorescent protein. Nat. Methods 20, 541–545 (2023).

51. St-Pierre, F. et al. One-Step Cloning and Chromosomal Integration of DNA. ACS Synth. Biol. 2, 537–541 (2013).

52. Guarino, A., Fiore, D., Salzano, D. & di Bernardo, M. Balancing Cell Populations Endowed with a Synthetic Toggle Switch via Adaptive Pulsatile Feedback Control. ACS Synth. Biol. 9, 793–803 (2020).

53. Salzano, D., Fiore, D. & di Bernardo, M. Controlling Reversible Cell Differentiation for Labor Division in Microbial Consortia. http://biorxiv.org/lookup/doi/10.1101/2021.08.03.454926 (2021) xdoi:10.1101/2021.08.03.454926.

54. Grob, A., Di Blasi, R. & Ceroni, F. Experimental tools to reduce the burden of bacterial synthetic biology. Curr. Opin. Syst. Biol. 28, 100393 (2021).

55. Sechkar, K., Steel, H., Perrino, G. & Stan, G.-B. A coarse-grained bacterial cell model for resource-aware analysis and design of synthetic gene circuits. Nat. Commun. 15, 1981 (2024).

56. Liu, Q., Schumacher, J., Wan, X., Lou, C. & Wang, B. Orthogonality and Burdens of Heterologous AND Gate Gene Circuits in E. coli. ACS Synth. Biol. 7, 553–564 (2018).

57. Zhang, Q., Azarin, S. M. & Sarkar, C. A. Model-guided engineering of DNA sequences with predictable site-specific recombination rates. Nat. Commun. 13, 4152 (2022).

58. Farruggio, A. P., Chavez, C. L., Mikell, C. L. & Calos, M. P. Efficient reversal of phiC31 integrase recombination in mammalian cells. Biotechnol. J. 7, 1332–1336 (2012).

59. Larsson, E. M., Murray, R. M. & Newman, D. K. Engineering the Soil Bacterium Pseudomonas synxantha 2–79 into a Ratiometric Bioreporter for Phosphorus Limitation. ACS Synth. Biol. 13, 384–393 (2024).

60. Mimee, M., Tucker, A. C., Voigt, C. A. & Lu, T. K. Programming a Human Commensal Bacterium, Bacteroides thetaiotaomicron, to Sense and Respond to Stimuli in the Murine Gut Microbiota. Cell Syst. 1, 62–71 (2015).

61. Wang, F. et al. One-pot biocatalytic route from cycloalkanes to α,ω-dicarboxylic acids by designed Escherichia coli consortia. Nat. Commun. 11, 5035 (2020).

62. Wang, X. et al. Engineering a Microbial Consortium Based Whole-Cell System for Efficient Production of Glutarate From L-Lysine. Front. Microbiol. 10, 341 (2019).

63. Zhang, Z. et al. One-pot biosynthesis of 1,6-hexanediol from cyclohexane by de novo designed cascade biocatalysis. Green Chem. 22, 7476–7483 (2020).

64. Nazir, A., Hussain, F. H. N. & Raza, A. Advancing microbiota therapeutics: the role of synthetic biology in engineering microbial communities for precision medicine. Front. Bioeng. Biotechnol. 12, 1511149 (2024).

65. An, B. et al. Programming Living Glue Systems to Perform Autonomous Mechanical Repairs. Matter 3, 2080–2092 (2020).

66. Gilbert, C. & Ellis, T. Biological Engineered Living Materials: Growing Functional Materials with Genetically Programmable Properties. ACS Synth. Biol. 8, 1–15 (2019).

