## Supplementary material for "Synthetic Phase Variation for Engineered Microbial Consortia": Main supplement (figures, tables and modeling)

June 11, 2025

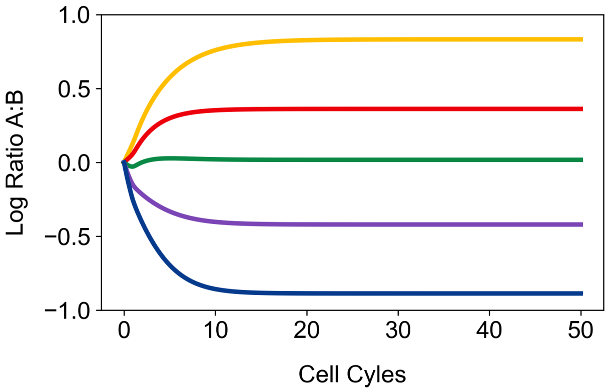


Figure S1: Composition tuning in the 2-state chemostat model. Different color traces correspond to different switching parameter sets ($\beta_j$ values) in the 2-state chemostat model, leading to different steady state compositions. Parameter sets used for this plot are available in the modeling section of the supplement.

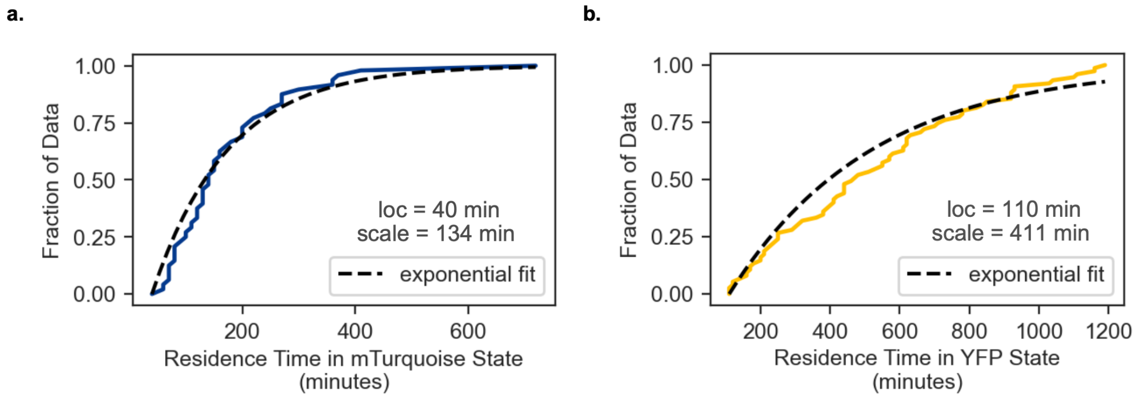


Figure S2: Mother machine residence time cumulative distribution fits. **a.** mTurquoise residence time cumulative distribution fit. **b.** YFP residence cumulative distribution time fit. Collected data was fit to an exponential distribution, allowing both the location and scale parameter to vary.

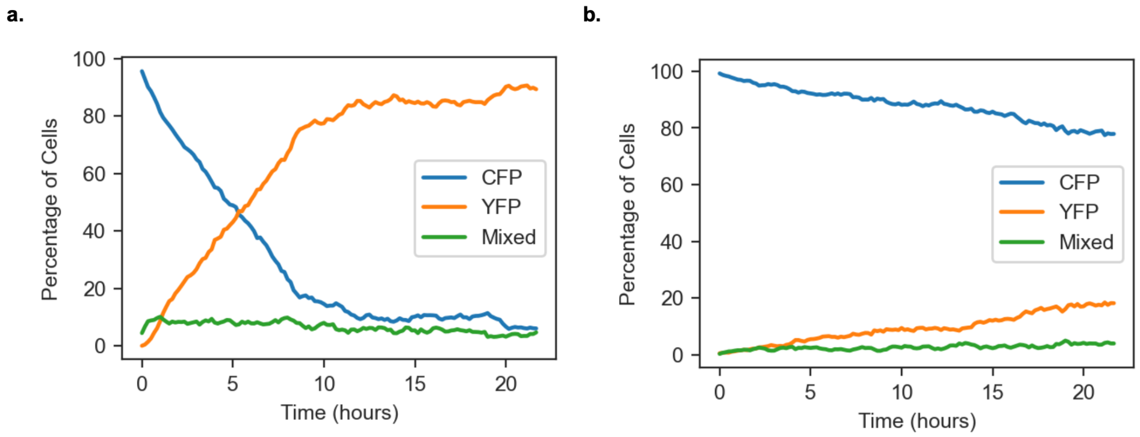


Figure S3: Mother machine individual population dynamics for 2-state system. **a.** YFP-dominant condition. **b.** mTurquoise-dominant condition. Cells are classified into YFP, mTurquoise or mixed bins according to their YFP to mTurquoise ratio. Cells abundance are expressed as percentage of live, fluorescent cells.

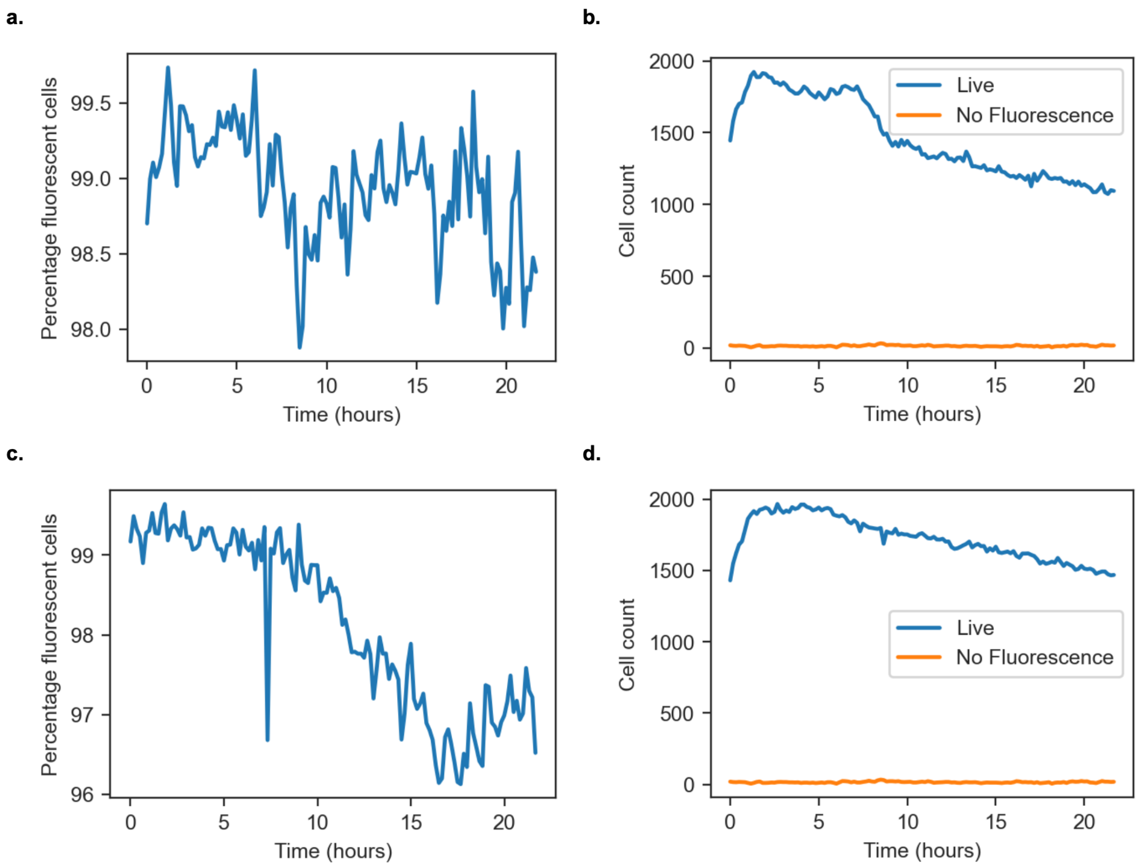


Figure S4: Mother machine cell viability dynamics for 2-state system. **a** YFP-dominant condition percentage fluorescent cell dynamics. **b**. YFP-dominant condition live and non-fluorescent cell abundance dynamics. **c** mTurquoise-dominant condition percentage fluorescent cell dynamics. **d**. mTurquoise-dominant condition live and non-fluorescent cell abundance dynamics. Cells are considered non-fluorescent cell when the sum of the normalized flourscent intensity is below 0.25.

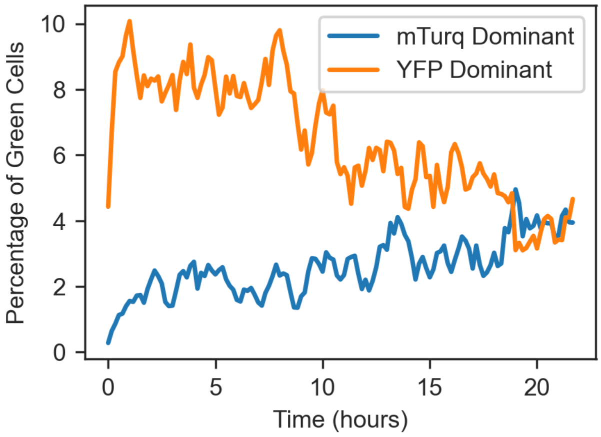


Figure S5: Mother machine mixed cell abundance dynamics. Cells are classified into YFP, mTurquoise or mixed bins according to their YFP to mTurquoise ratio. Cells abundance are expressed as percentage of live, fluorescent cells.

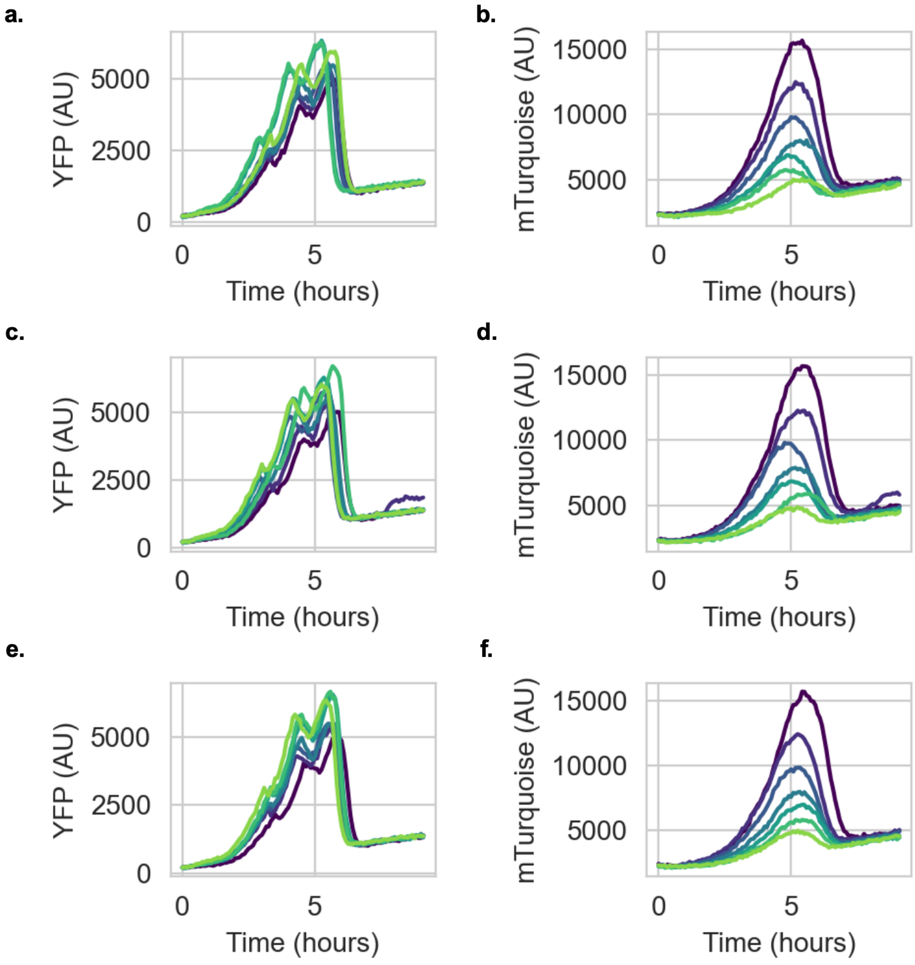


Figure S6: Raw fluorescent traces for 2-strain uncontrolled consortia. **a.** YFP dynamics for replicate 1. **b.** mTurquoise dynamics for replicate 1. **c.** YFP dynamics for replicate 2. **d.** mTurquoise dynamics for replicate 2. **e.** YFP dynamics for replicate 3. **f.** mTurquoise dynamics for replicate 3. Each individual peak corresponds to a growth cycle for a given well. Early cycles are plotted in dark colors, starting in dark blue and progress to light colors as cycle number increases, finishing in light green.

SO 1

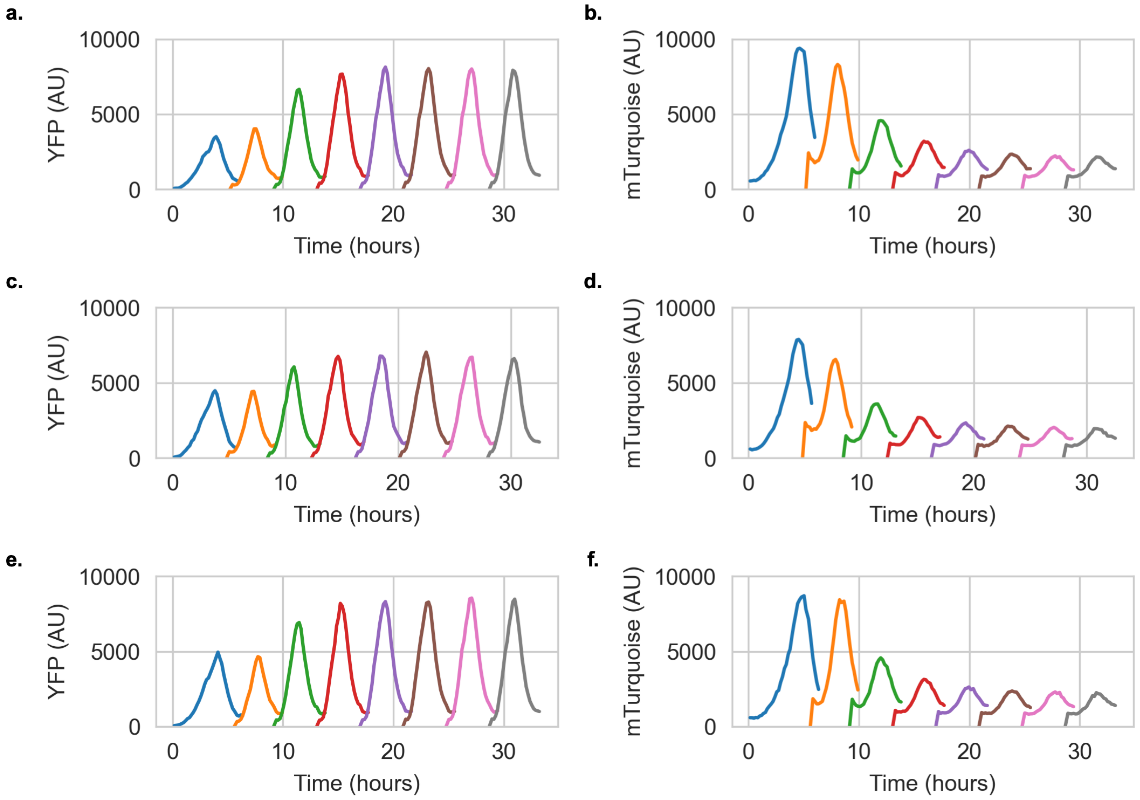


Figure S7: Raw 2-state composition control fluorescent traces for condition A ▬. **a.** YFP dynamics for initial condition 1. **b.** mTurquoise dynamics for initial condition 1. **c.** YFP dynamics for initial condition 2. **d.** mTurquoise dynamics for initial condition 2. **e.** YFP dynamics for initial condition 3. **f.** mTurquoise dynamics for initial condition 3. Each individual peak corresponds to a growth cycle for a given well, max fluorescent signal during this growth signal was taken to be representative of composition for a given fluorescent channel.

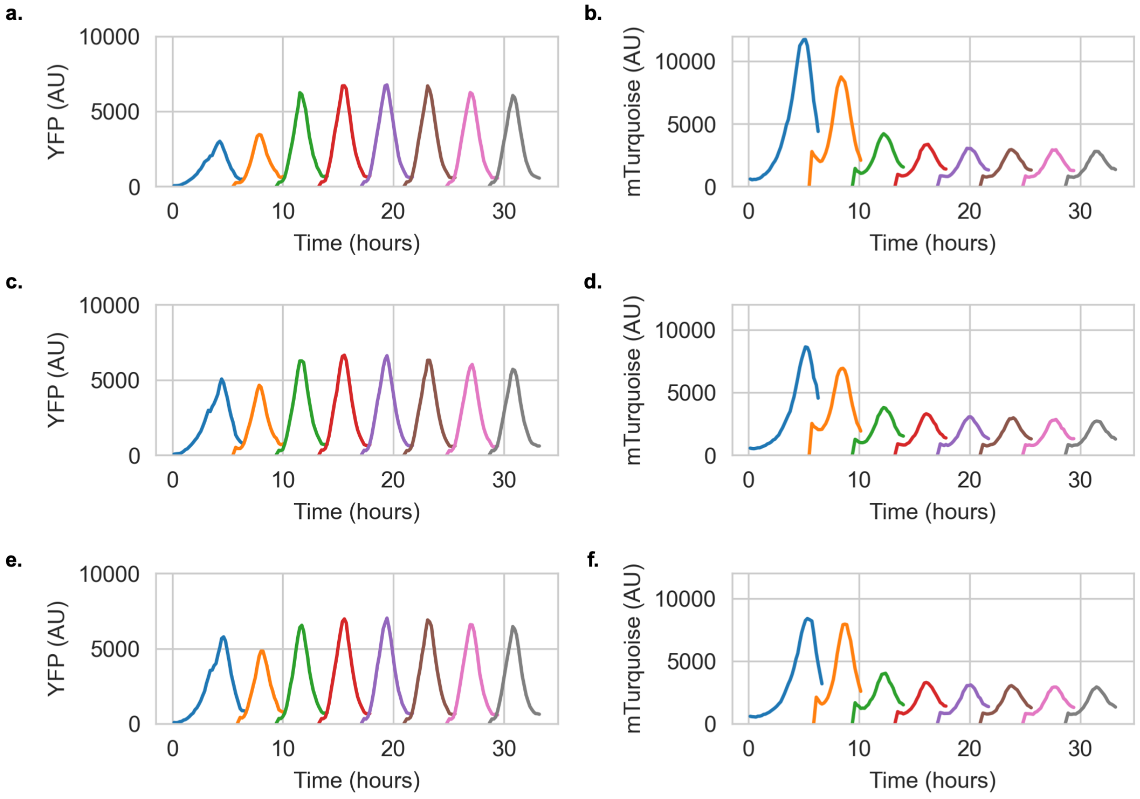


Figure S8: Raw 2-state composition control fluorescent traces for condition B . **a.** YFP dynamics for initial condition 1. **b.** mTurquoise dynamics for initial condition 1. **c.** YFP dynamics for initial condition 2. **d.** mTurquoise dynamics for initial condition 2. **e.** YFP dynamics for initial condition 3. **f.** mTurquoise dynamics for initial condition 3. Each individual peak corresponds to a growth cycle for a given well, max fluorescent signal during this growth signal was taken to be representative of composition for a given fluorescent channel.

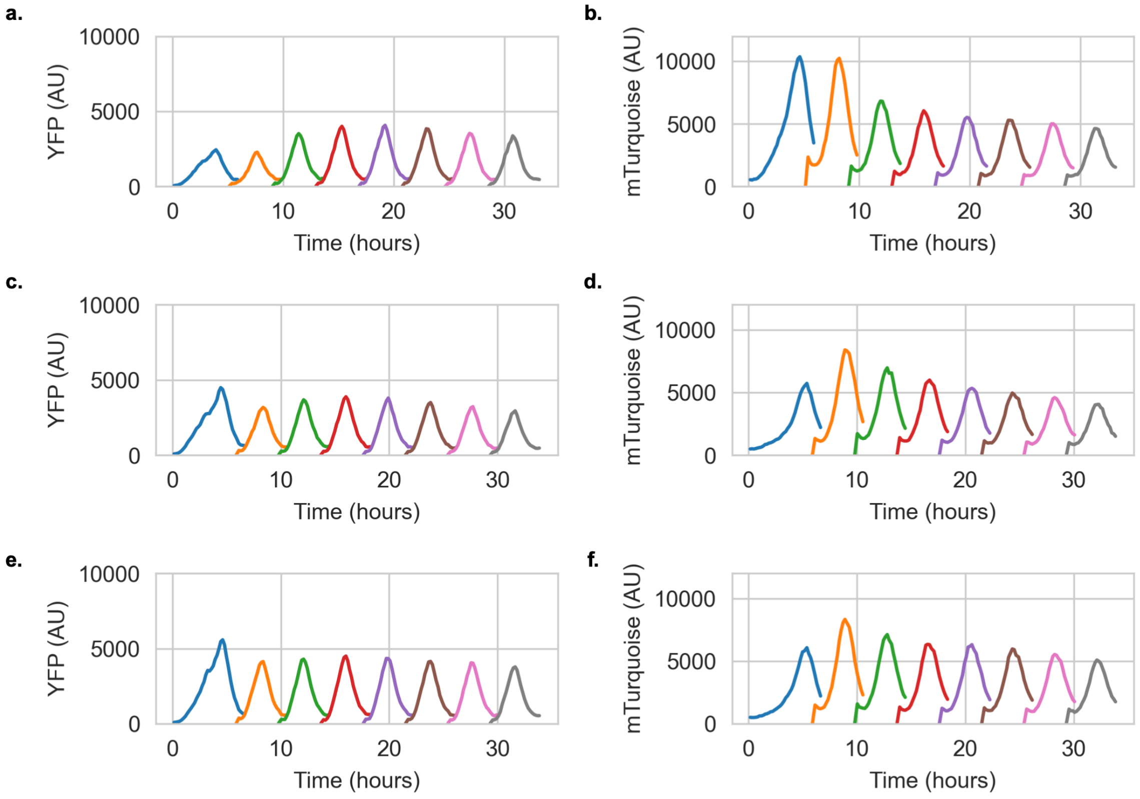


Figure S9: Raw 2-state composition control fluorescent traces for condition C ▬. **a.** YFP dynamics for initial condition 1. **b.** mTurquoise dynamics for initial condition 1. **c.** YFP dynamics for initial condition 2. **d.** mTurquoise dynamics for initial condition 2. **e.** YFP dynamics for initial condition 3. **f.** mTurquoise dynamics for initial condition 3. Each individual peak corresponds to a growth cycle for a given well, max fluorescent signal during this growth signal was taken to be representative of composition for a given fluorescent channel.

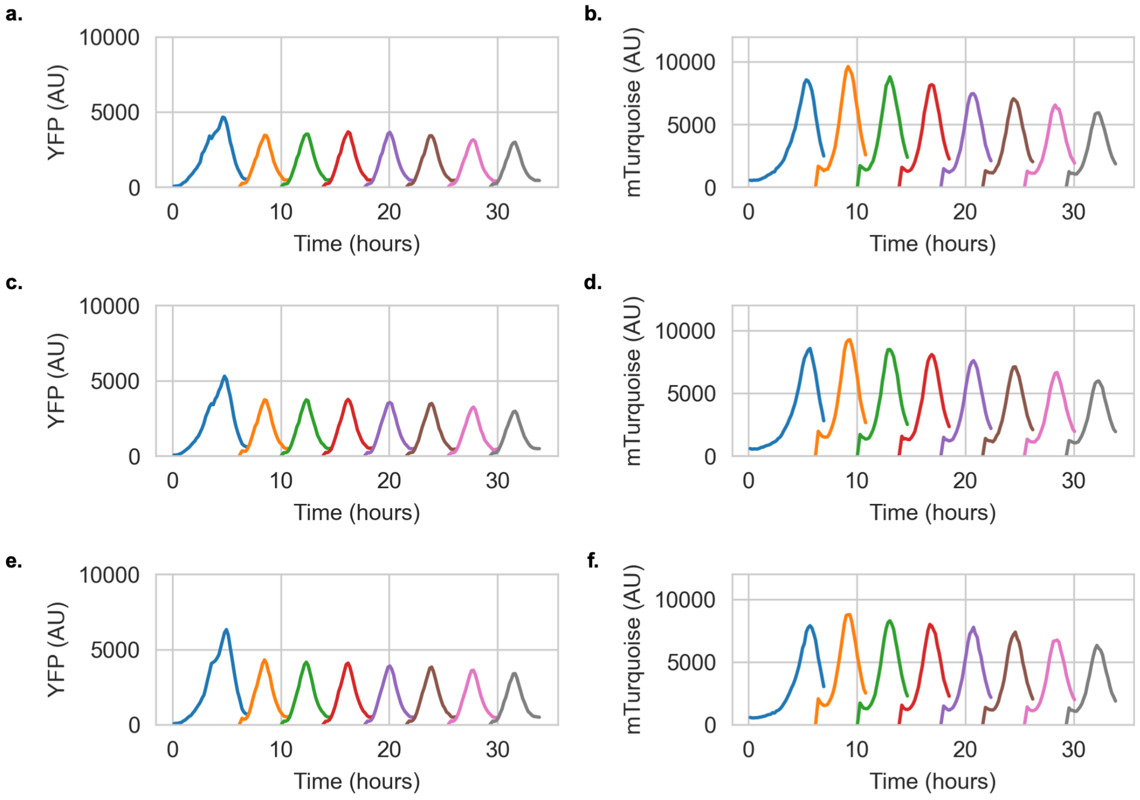


Figure S10: Raw 2-state composition control fluorescent traces for condition D. **a.** YFP dynamics for initial condition 1. **b.** mTurquoise dynamics for initial condition 1. **c.** YFP dynamics for initial condition 2. **d.** mTurquoise dynamics for initial condition 2. **e.** YFP dynamics for initial condition 3. **f.** mTurquoise dynamics for initial condition 3. Each individual peak corresponds to a growth cycle for a given well, max fluorescent signal during this growth signal was taken to be representative of composition for a given fluorescent channel.

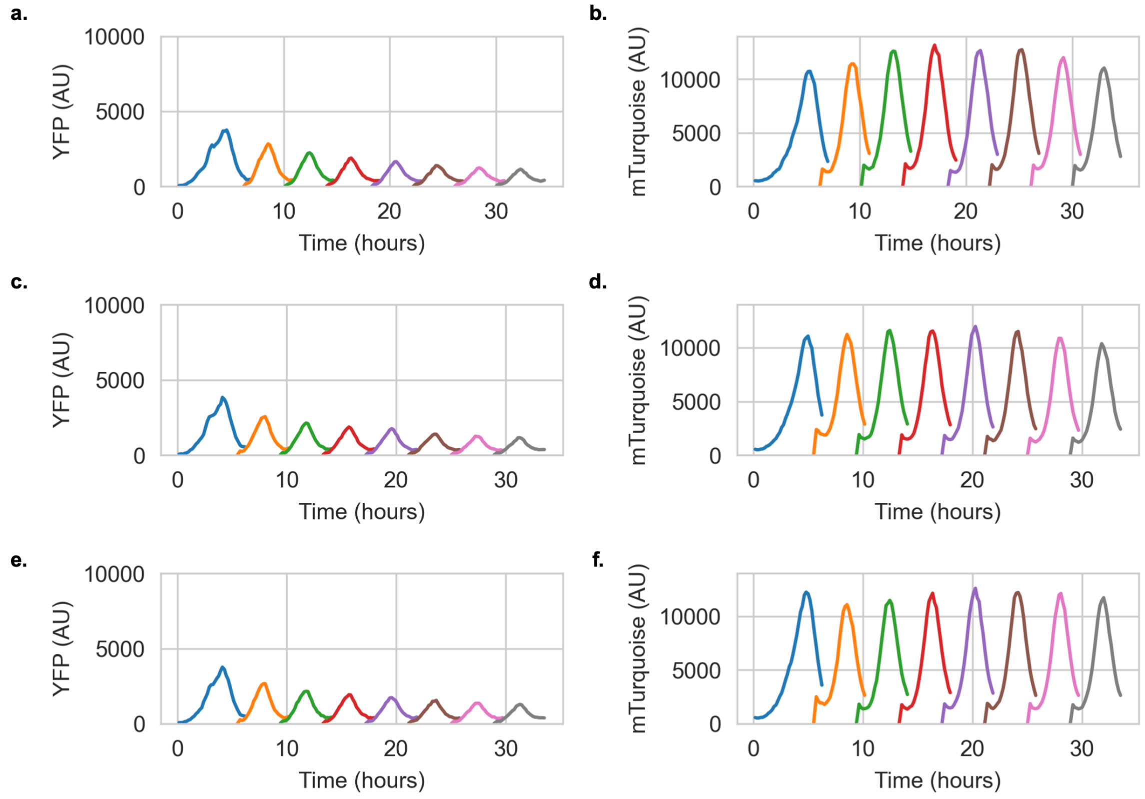


Figure S11: Raw 2-state composition control fluorescent traces for condition E ▬. **a.** YFP dynamics for initial condition 1. **b.** mTurquoise dynamics for initial condition 1. **c.** YFP dynamics for initial condition 2. **d.** mTurquoise dynamics for initial condition 2. **e.** YFP dynamics for initial condition 3. **f.** mTurquoise dynamics for initial condition 3. Each individual peak corresponds to a growth cycle for a given well, max fluorescent signal during this growth signal was taken to be representative of composition for a given fluorescent channel.

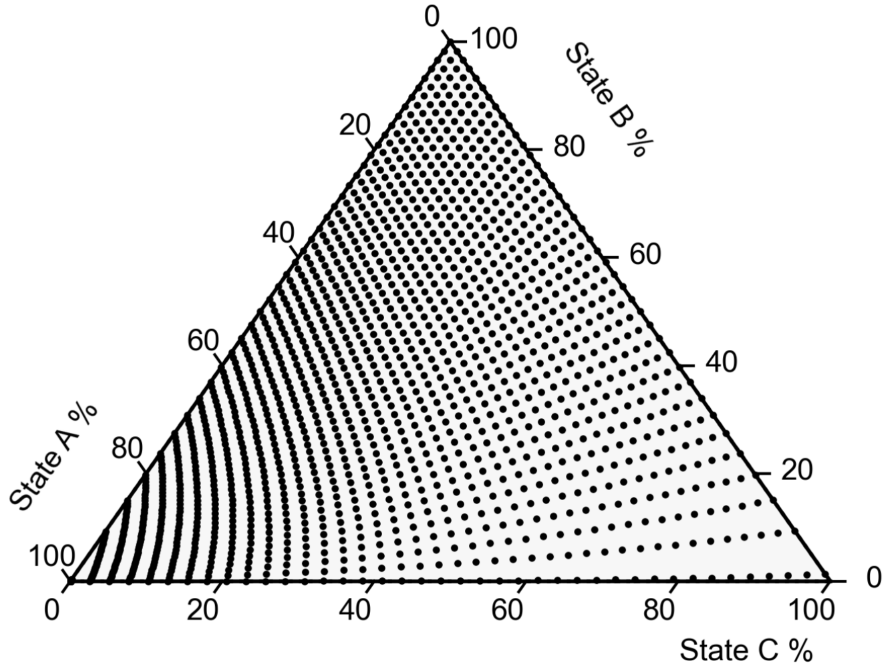


Figure S12: Parameter sweeps of the 3-state model shows our approach can occupy any position in composition space . Ternary diagram of the 3-state system, each dot corresponds to the steady-state composition for a given set of switching rates (four distinct $\beta_j$ values). The parameter ranges used for this plot are available in the modeling section of the supplement.

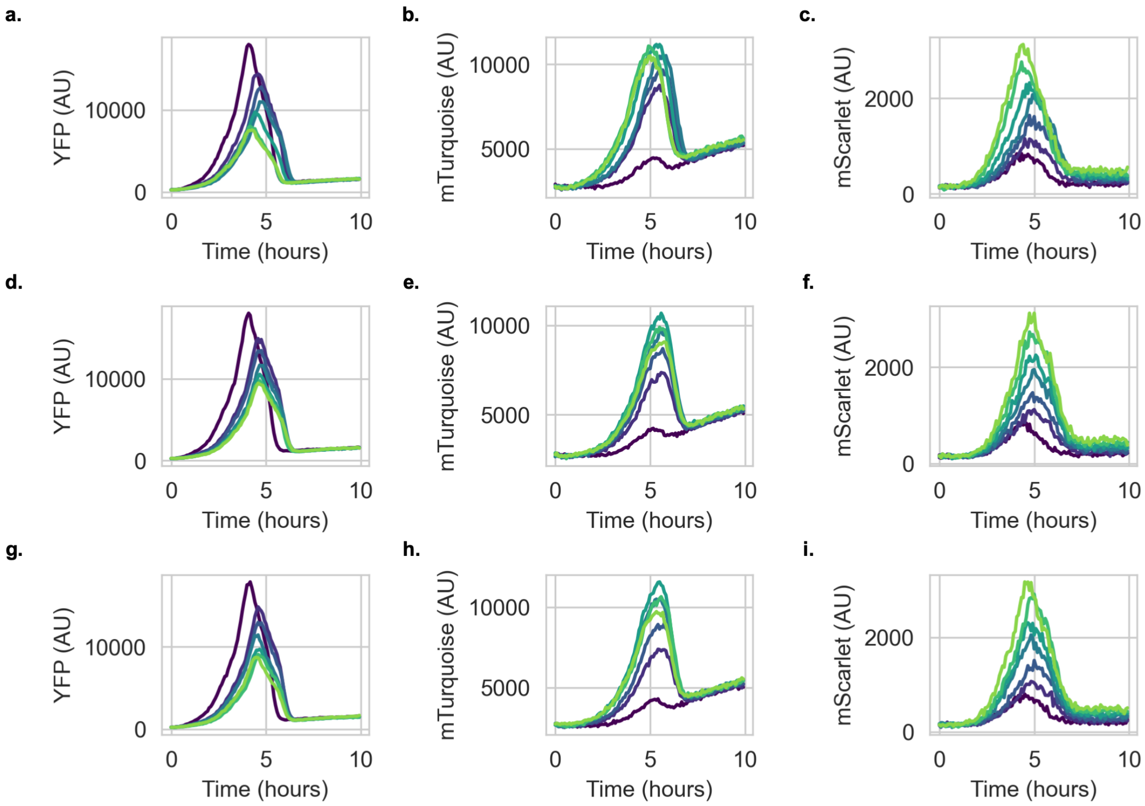


Figure S13: Raw 3-state composition control fluorescent traces for condition A. **a.** YFP dynamics for replicate 1. **b.** mTurquoise dynamics for replicate 1. **c.** mScarlet dynamics for replicate 1. **d.** YFP dynamics for replicate 2. **e.** mTurquoise dynamics for replicate 2. **f.** mScarlet dynamics for replicate 2. **g.** YFP dynamics for replicate 3. **h.** mTurquoise dynamics for replicate 3. **i.** mScarlet dynamics for replicate 3. Each individual peak corresponds to a growth cycle for a given well. Early cycles are plotted in dark colors, starting in dark blue and progress to light colors as cycle number increases, finishing in light green. Max fluorescent signal during this growth signal was taken to be representative of composition for a given fluorescent channel.

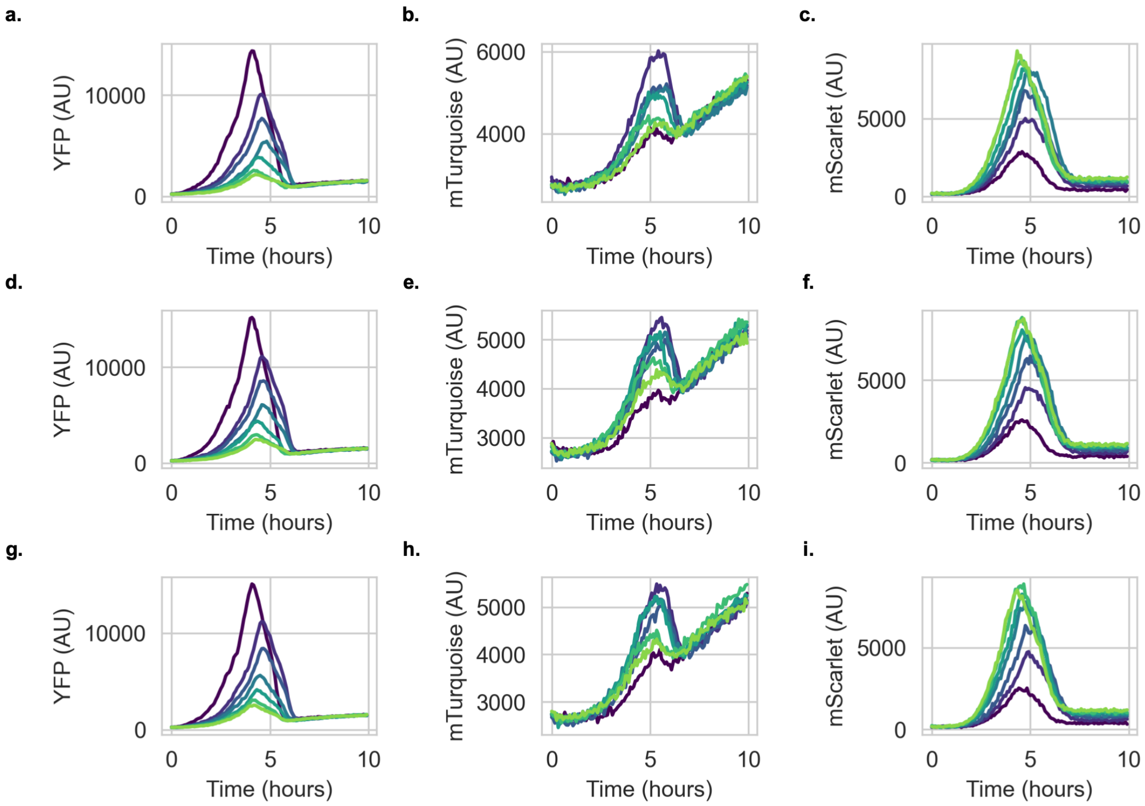


Figure S14: Raw 3-state composition control fluorescent traces for condition B. **a.** YFP dynamics for replicate 1. **b.** mTurquoise dynamics for replicate 1. **c.** mScarlet dynamics for replicate 1. **d.** YFP dynamics for replicate 2. **e.** mTurquoise dynamics for replicate 2. **f.** mScarlet dynamics for replicate 2. **g.** YFP dynamics for replicate 3. **h.** mTurquoise dynamics for replicate 3. **i.** mScarlet dynamics for replicate 3. Each individual peak corresponds to a growth cycle for a given well. Early cycles are plotted in dark colors, starting in dark blue and progress to light colors as cycle number increases, finishing in light green. Max fluorescent signal during this growth signal was taken to be representative of composition for a given fluorescent channel.

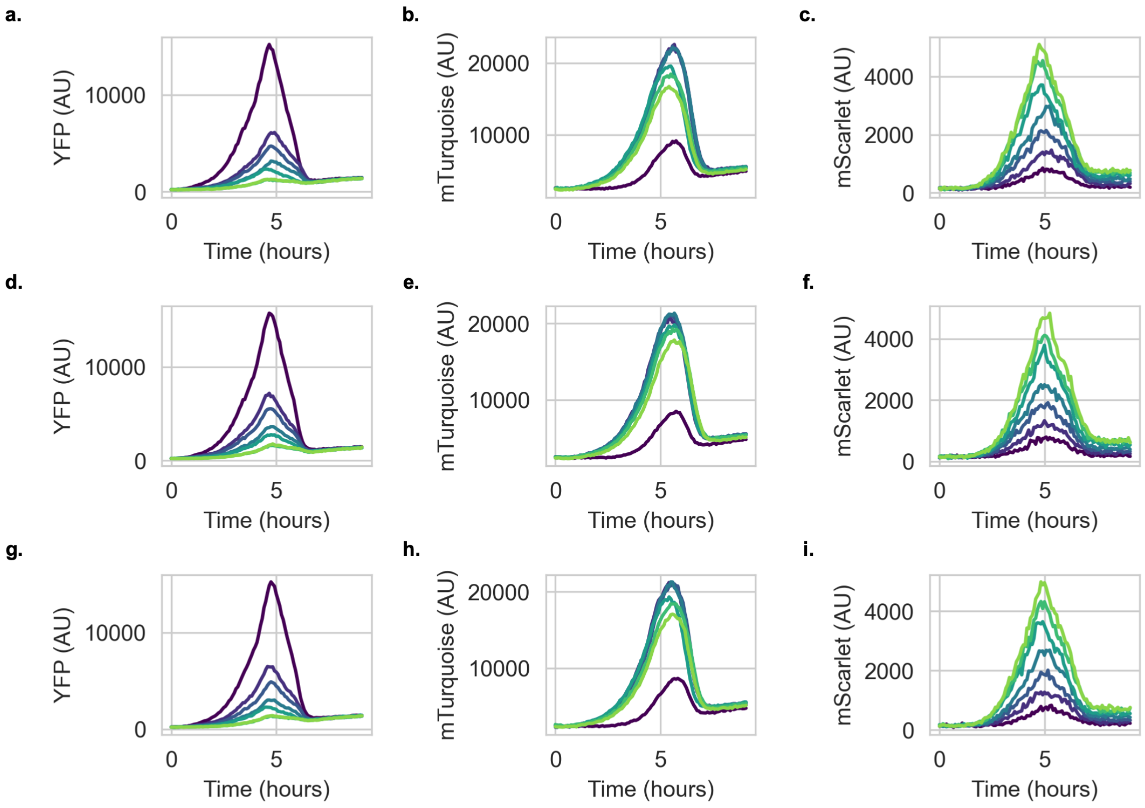


Figure S15: Raw 3-state composition control fluorescent traces for condition C. **a.** YFP dynamics for replicate 1. **b.** mTurquoise dynamics for replicate 1. **c.** mScarlet dynamics for replicate 1. **d.** YFP dynamics for replicate 2. **e.** mTurquoise dynamics for replicate 2. **f.** mScarlet dynamics for replicate 2. **g.** YFP dynamics for replicate 3. **h.** mTurquoise dynamics for replicate 3. **i.** mScarlet dynamics for replicate 3. Each individual peak corresponds to a growth cycle for a given well. Early cycles are plotted in dark colors, starting in dark blue and progress to light colors as cycle number increases, finishing in light green. Max fluorescent signal during this growth signal was taken to be representative of composition for a given fluorescent channel.

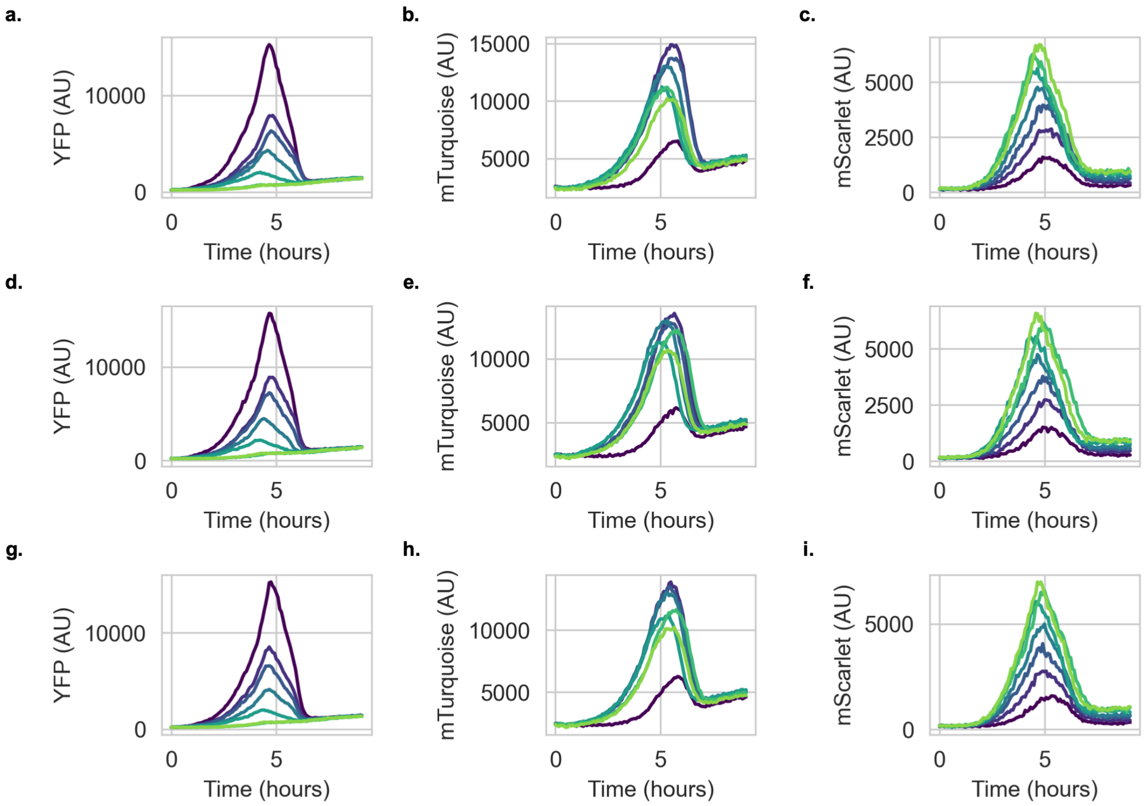


Figure S16: Raw 3-state composition control fluorescent traces for condition D. **a.** YFP dynamics for replicate 1. **b.** mTurquoise dynamics for replicate 1. **c.** mScarlet dynamics for replicate 1. **d.** YFP dynamics for replicate 2. **e.** mTurquoise dynamics for replicate 2. **f.** mScarlet dynamics for replicate 2. **g.** YFP dynamics for replicate 3. **h.** mTurquoise dynamics for replicate 3. **i.** mScarlet dynamics for replicate 3. Each individual peak corresponds to a growth cycle for a given well. Early cycles are plotted in dark colors, starting in dark blue and progress to light colors as cycle number increases, finishing in light green. Max fluorescent signal during this growth signal was taken to be representative of composition for a given fluorescent channel.

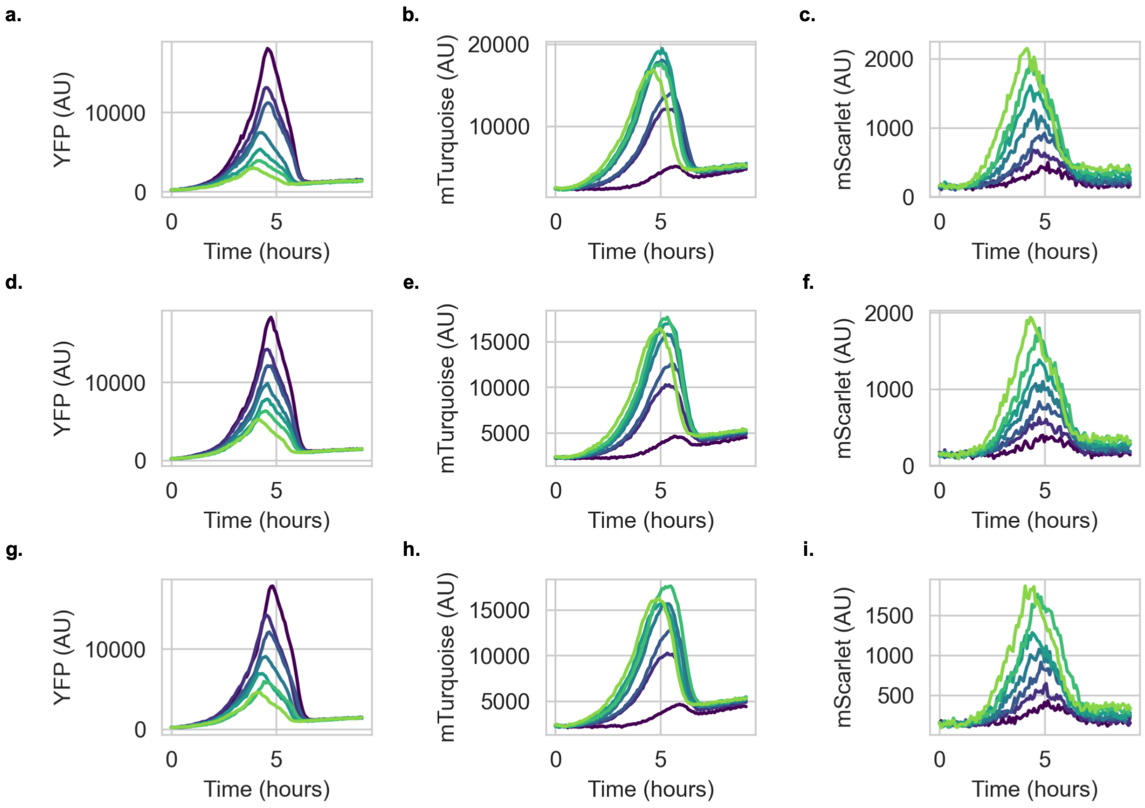


Figure S17: Raw 3-state composition control fluorescent traces for condition E. **a.** YFP dynamics for replicate 1. **b.** mTurquoise dynamics for replicate 1. **c.** mScarlet dynamics for replicate 1. **d.** YFP dynamics for replicate 2. **e.** mTurquoise dynamics for replicate 2. **f.** mScarlet dynamics for replicate 2. **g.** YFP dynamics for replicate 3. **h.** mTurquoise dynamics for replicate 3. **i.** mScarlet dynamics for replicate 3. Each individual peak corresponds to a growth cycle for a given well. Early cycles are plotted in dark colors, starting in dark blue and progress to light colors as cycle number increases, finishing in light green. Max fluorescent signal during this growth signal was taken to be representative of composition for a given fluorescent channel.

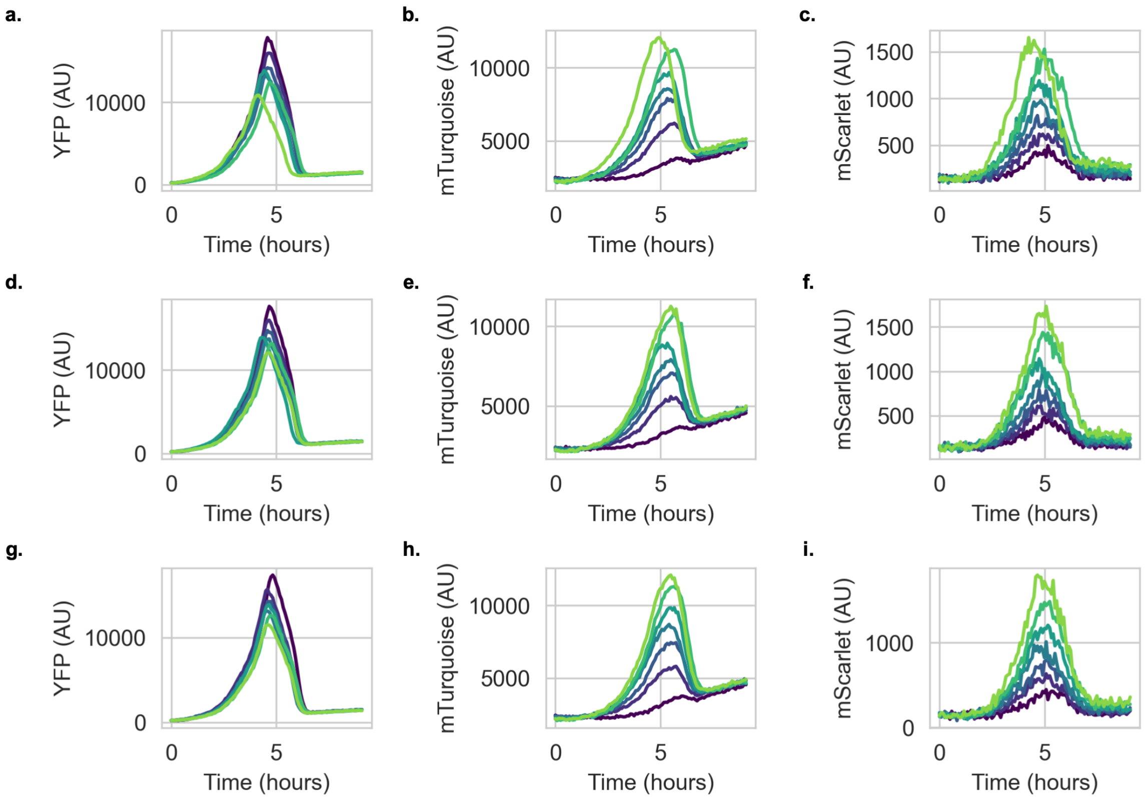


Figure S18: Raw 3-state composition control fluorescent traces for condition F. **a.** YFP dynamics for replicate 1. **b.** mTurquoise dynamics for replicate 1. **c.** mScarlet dynamics for replicate 1. **d.** YFP dynamics for replicate 2. **e.** mTurquoise dynamics for replicate 2. **f.** mScarlet dynamics for replicate 2. **g.** YFP dynamics for replicate 3. **h.** mTurquoise dynamics for replicate 3. **i.** mScarlet dynamics for replicate 3. Each individual peak corresponds to a growth cycle for a given well. Early cycles are plotted in dark colors, starting in dark blue and progress to light colors as cycle number increases, finishing in light green. Max fluorescent signal during this growth signal was taken to be representative of composition for a given fluorescent channel.

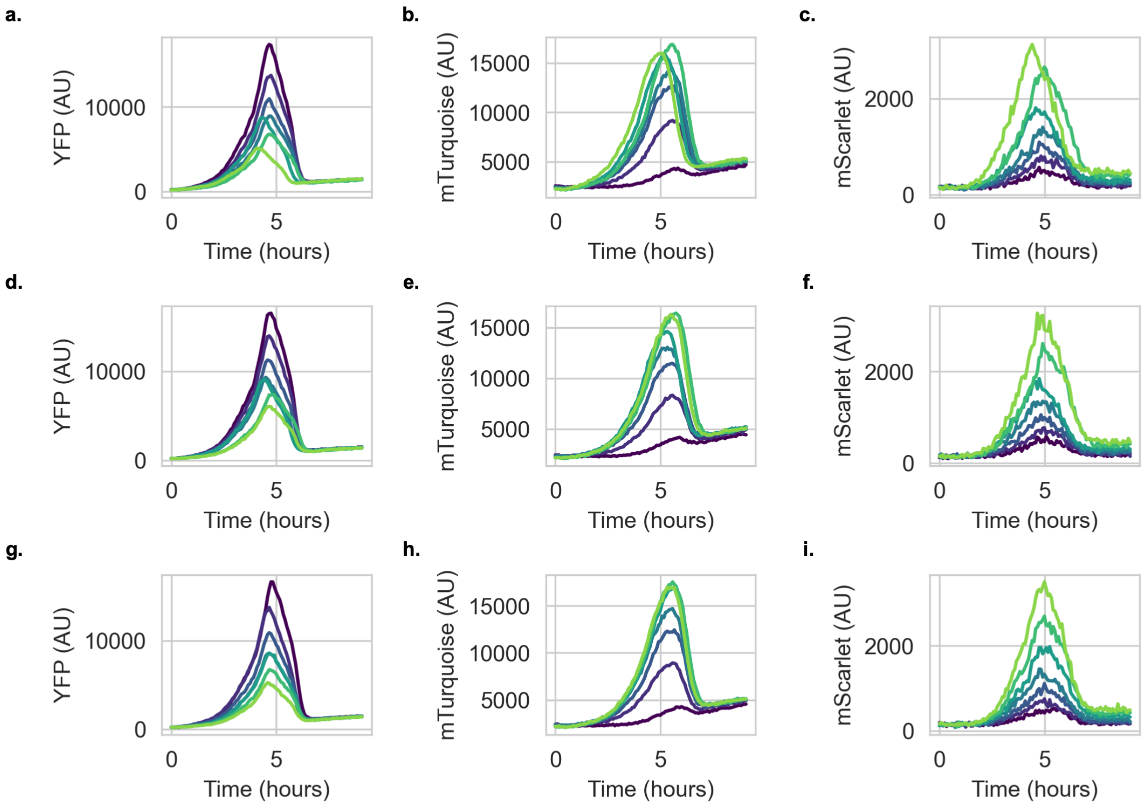


Figure S19: Raw 3-state composition control fluorescent traces for condition G. **a.** YFP dynamics for replicate 1. **b.** mTurquoise dynamics for replicate 1. **c.** mScarlet dynamics for replicate 1. **d.** YFP dynamics for replicate 2. **e.** mTurquoise dynamics for replicate 2. **f.** mScarlet dynamics for replicate 2. **g.** YFP dynamics for replicate 3. **h.** mTurquoise dynamics for replicate 3. **i.** mScarlet dynamics for replicate 3. Each individual peak corresponds to a growth cycle for a given well. Early cycles are plotted in dark colors, starting in dark blue and progress to light colors as cycle number increases, finishing in light green. Max fluorescent signal during this growth signal was taken to be representative of composition for a given fluorescent channel.

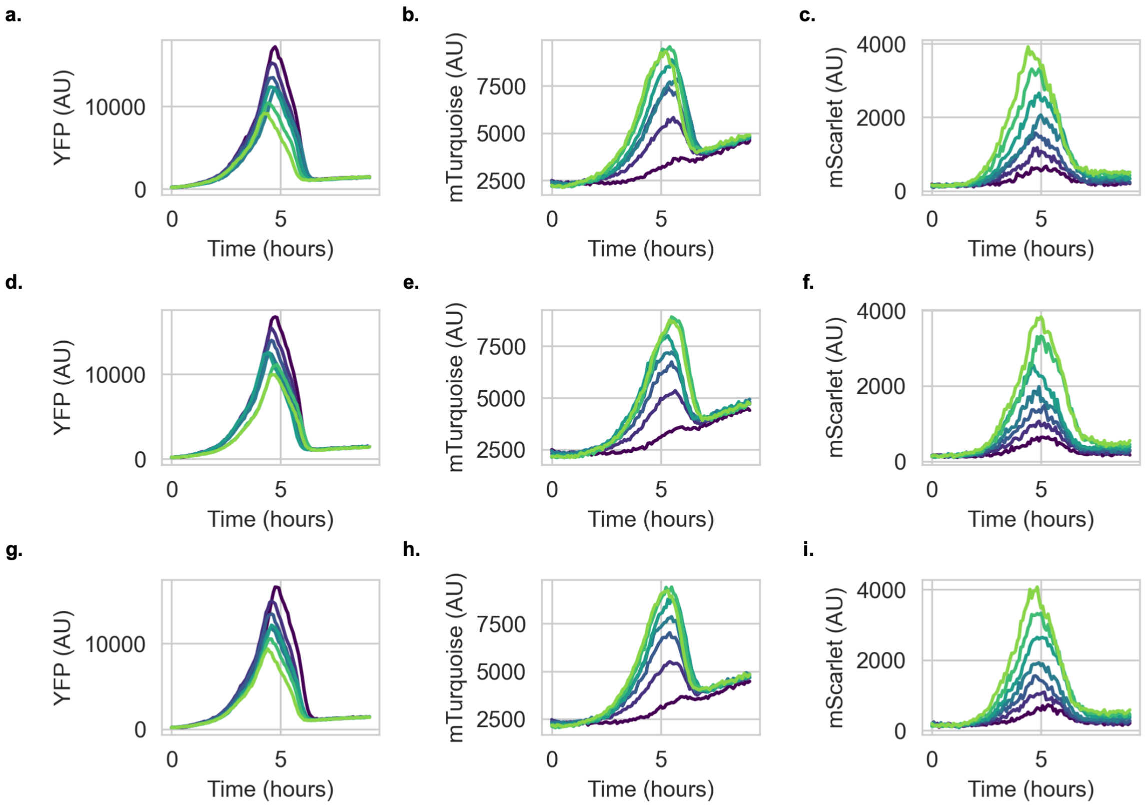


Figure S20: Raw 3-state composition control fluorescent traces for condition H. **a.** YFP dynamics for replicate 1. **b.** mTurquoise dynamics for replicate 1. **c.** mScarlet dynamics for replicate 1. **d.** YFP dynamics for replicate 2. **e.** mTurquoise dynamics for replicate 2. **f.** mScarlet dynamics for replicate 2. **g.** YFP dynamics for replicate 3. **h.** mTurquoise dynamics for replicate 3. **i.** mScarlet dynamics for replicate 3. Each individual peak corresponds to a growth cycle for a given well. Early cycles are plotted in dark colors, starting in dark blue and progress to light colors as cycle number increases, finishing in light green. Max fluorescent signal during this growth signal was taken to be representative of composition for a given fluorescent channel.

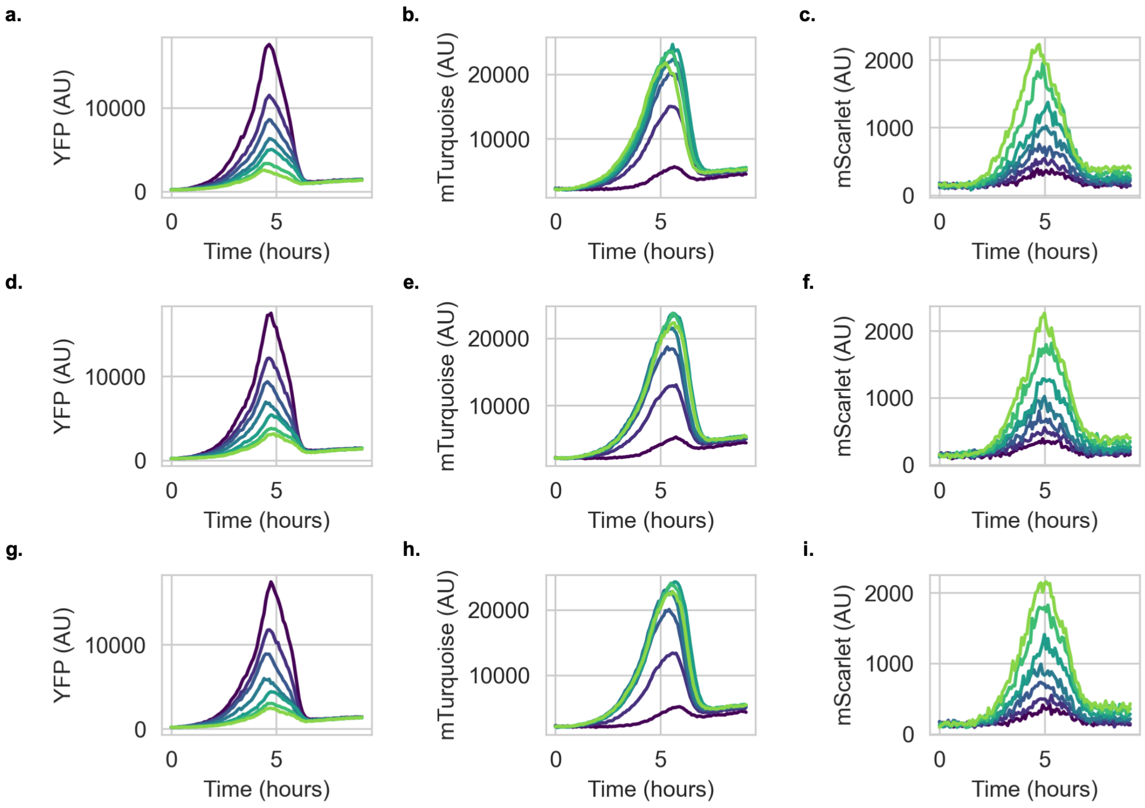


Figure S21: Raw 3-state composition control fluorescent traces for condition I. **a.** YFP dynamics for replicate 1. **b.** mTurquoise dynamics for replicate 1. **c.** mScarlet dynamics for replicate 1. **d.** YFP dynamics for replicate 2. **e.** mTurquoise dynamics for replicate 2. **f.** mScarlet dynamics for replicate 2. **g.** YFP dynamics for replicate 3. **h.** mTurquoise dynamics for replicate 3. **i.** mScarlet dynamics for replicate 3. Each individual peak corresponds to a growth cycle for a given well. Early cycles are plotted in dark colors, starting in dark blue and progress to light colors as cycle number increases, finishing in light green. Max fluorescent signal during this growth signal was taken to be representative of composition for a given fluorescent channel.

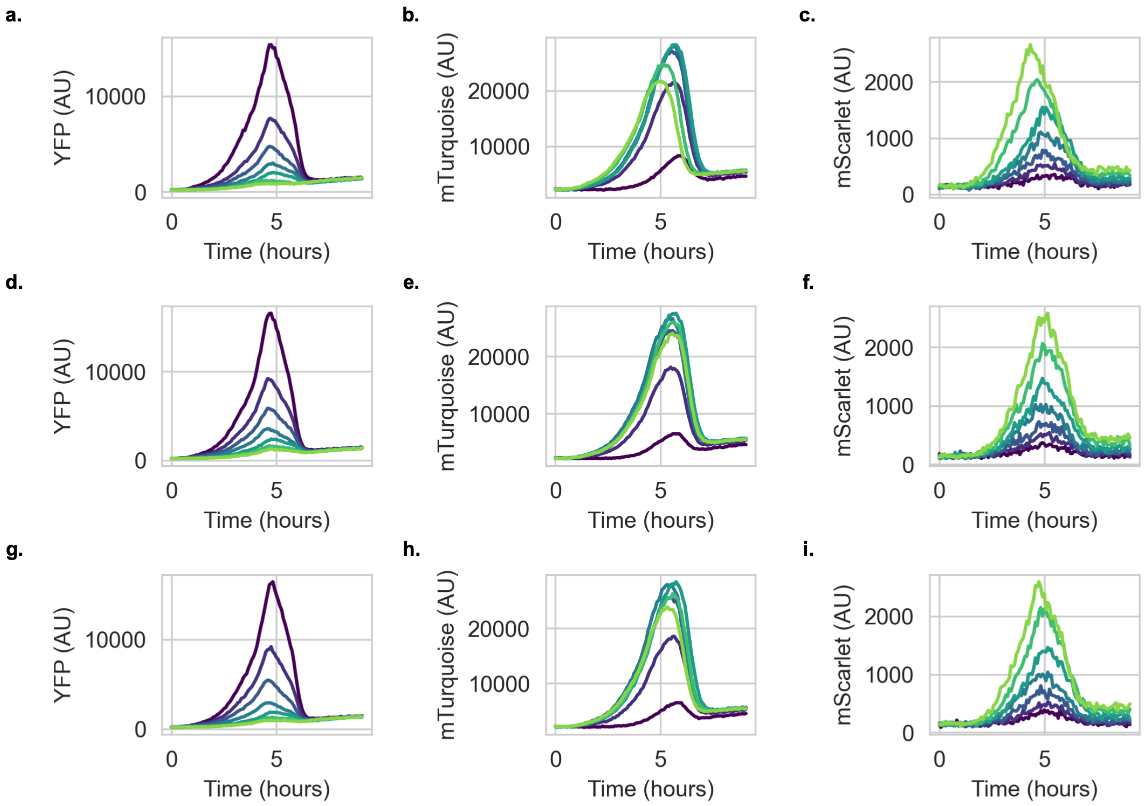


Figure S22: Raw 3-state composition control fluorescent traces for condition J. **a.** YFP dynamics for replicate 1. **b.** mTurquoise dynamics for replicate 1. **c.** mScarlet dynamics for replicate 1. **d.** YFP dynamics for replicate 2. **e.** mTurquoise dynamics for replicate 2. **f.** mScarlet dynamics for replicate 2. **g.** YFP dynamics for replicate 3. **h.** mTurquoise dynamics for replicate 3. **i.** mScarlet dynamics for replicate 3. Each individual peak corresponds to a growth cycle for a given well. Early cycles are plotted in dark colors, starting in dark blue and progress to light colors as cycle number increases, finishing in light green. Max fluorescent signal during this growth signal was taken to be representative of composition for a given fluorescent channel.

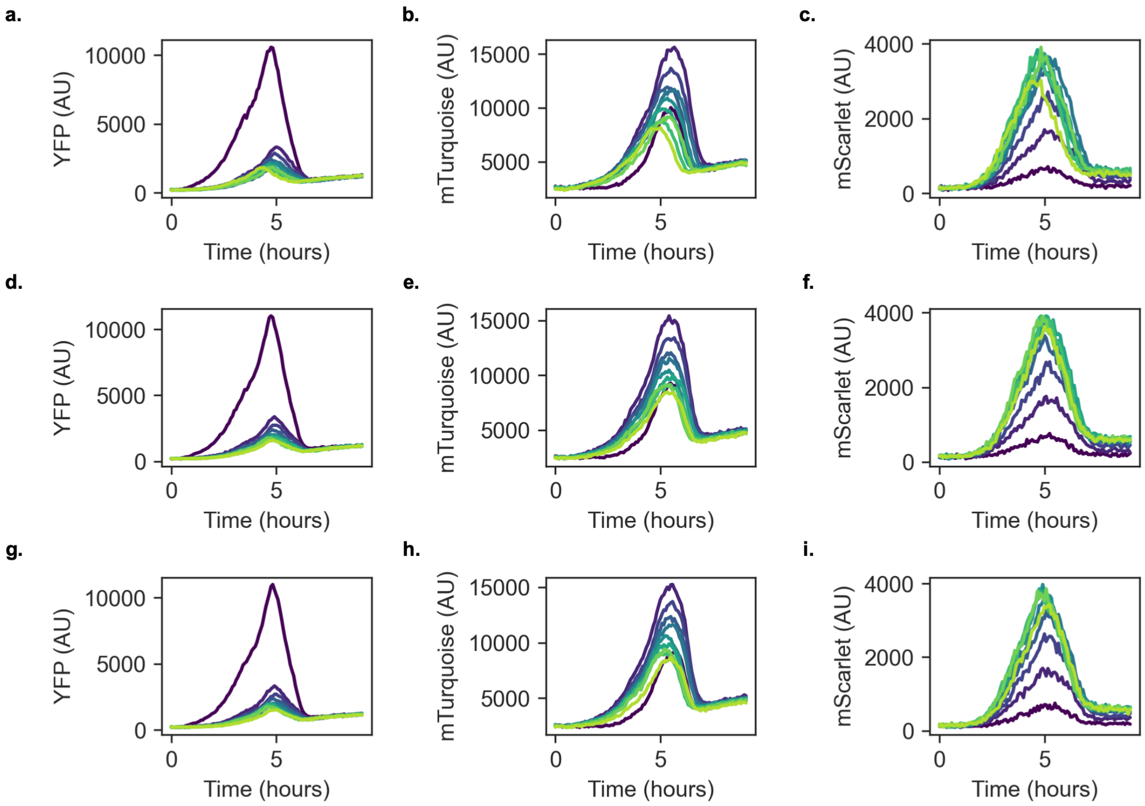


Figure S23: Raw 3-state composition control fluorescent traces for condition K. **a.** YFP dynamics for replicate 1. **b.** mTurquoise dynamics for replicate 1. **c.** mScarlet dynamics for replicate 1. **d.** YFP dynamics for replicate 2. **e.** mTurquoise dynamics for replicate 2. **f.** mScarlet dynamics for replicate 2. **g.** YFP dynamics for replicate 3. **h.** mTurquoise dynamics for replicate 3. **i.** mScarlet dynamics for replicate 3. Each individual peak corresponds to a growth cycle for a given well. Early cycles are plotted in dark colors, starting in dark blue and progress to light colors as cycle number increases, finishing in light green. Max fluorescent signal during this growth signal was taken to be representative of composition for a given fluorescent channel.

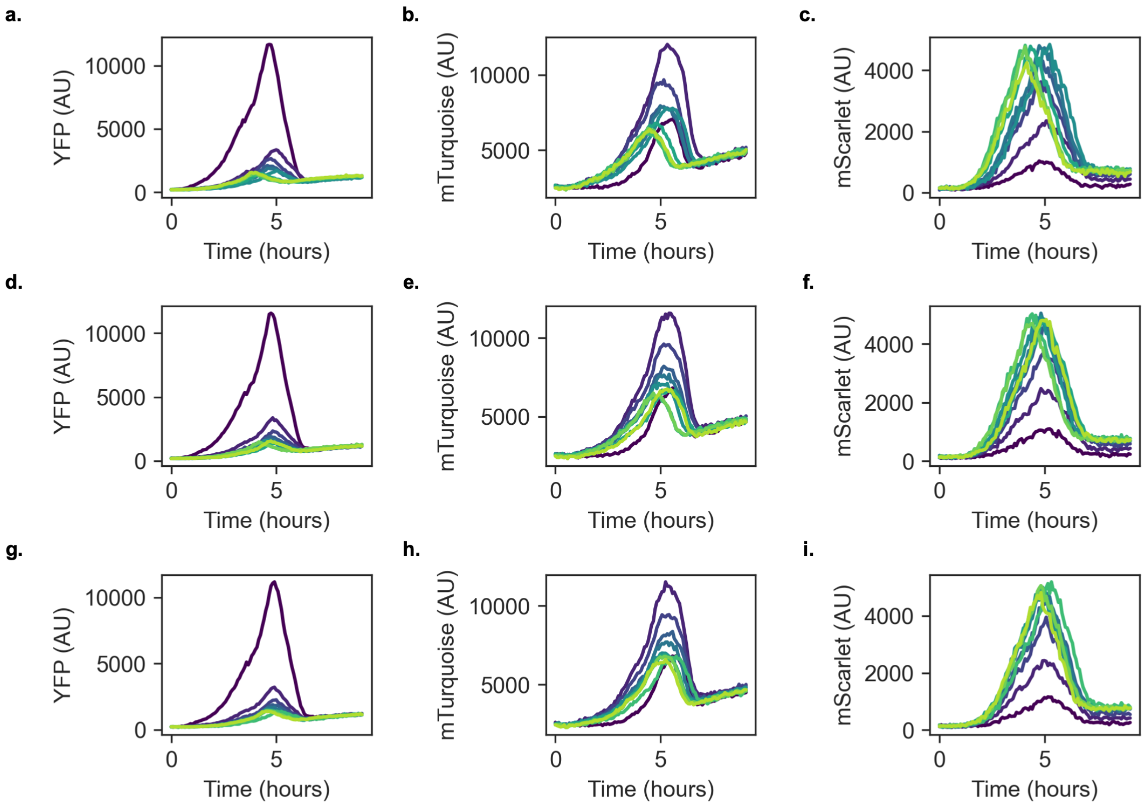


Figure S24: Raw 3-state composition control fluorescent traces for condition L. **a.** YFP dynamics for replicate 1. **b.** mTurquoise dynamics for replicate 1. **c.** mScarlet dynamics for replicate 1. **d.** YFP dynamics for replicate 2. **e.** mTurquoise dynamics for replicate 2. **f.** mScarlet dynamics for replicate 2. **g.** YFP dynamics for replicate 3. **h.** mTurquoise dynamics for replicate 3. **i.** mScarlet dynamics for replicate 3. Each individual peak corresponds to a growth cycle for a given well. Early cycles are plotted in dark colors, starting in dark blue and progress to light colors as cycle number increases, finishing in light green. Max fluorescent signal during this growth signal was taken to be representative of composition for a given fluorescent channel.

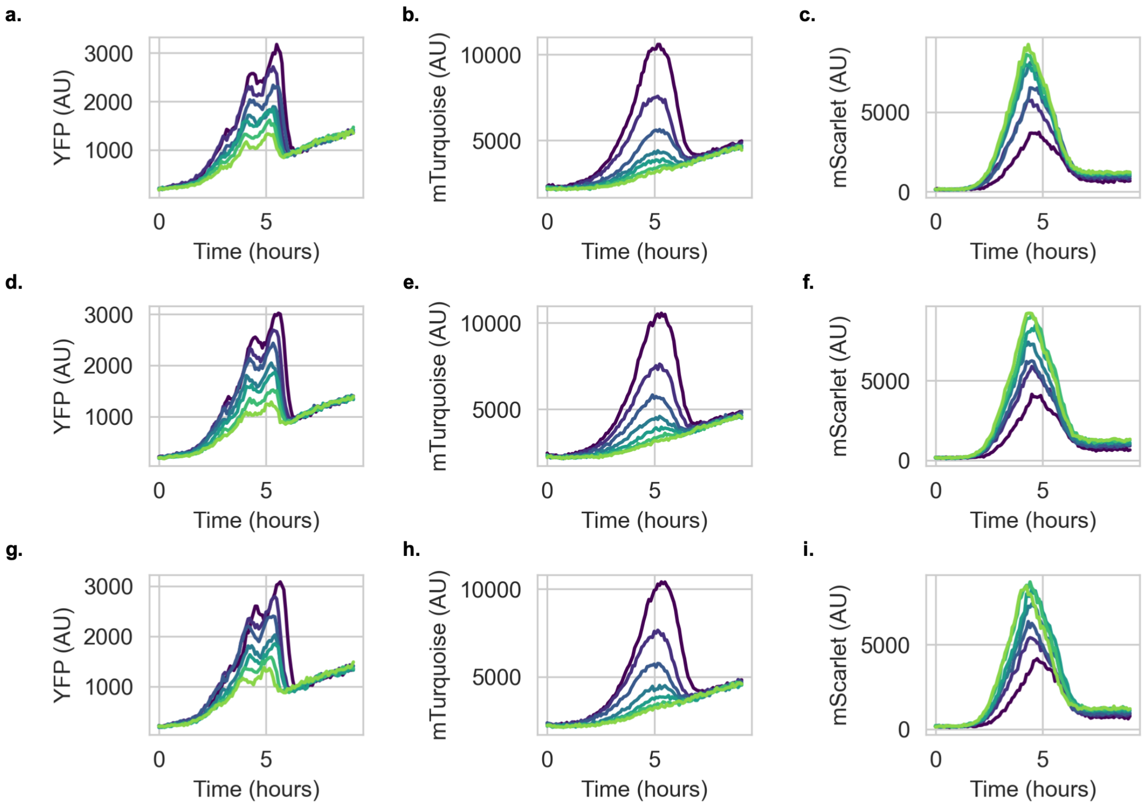


Figure S25: Raw fluorescent traces for 3-strain uncontrolled consortia. **a.** YFP dynamics for replicate 1. **b.** mTurquoise dynamics for replicate 1. **c.** mScarlet dynamics for replicate 1. **d.** YFP dynamics for replicate 2. **e.** mTurquoise dynamics for replicate 2. **f.** mScarlet dynamics for replicate 2. **g.** YFP dynamics for replicate 3. **h.** mTurquoise dynamics for replicate 3. **i.** mScarlet dynamics for replicate 3. Each individual peak corresponds to a growth cycle for a given well. Early cycles are plotted in dark colors, starting in dark blue and progress to light colors as cycle number increases, finishing in light green. Max fluorescent signal during this growth signal was taken to be representative of composition for a given fluorescent channel.

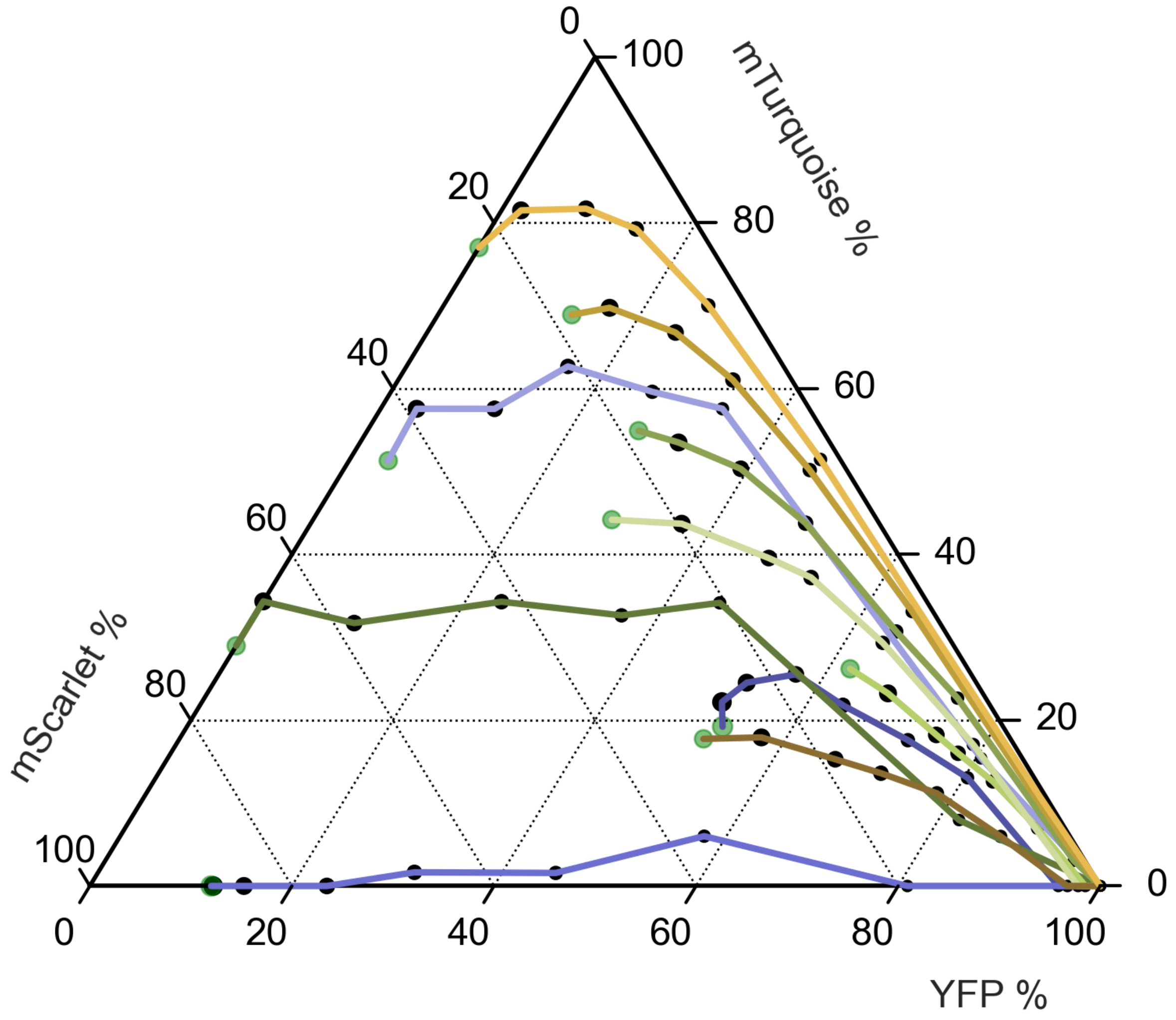


Figure S26: 3-state composition control traces. Different color traces correspond to different inducer conditions described in Table S1. Fluorescence values have been normalized to the max possible fluorescence signal for the given channel. As typical for a ternary diagram, the ratio of all three channels is used to calculate positions of individual trajectories. Each point in a given trajectory is separated by a fixed growth cycle of 12 hours.

| Condition | | Salicylate ($\mu M$) | IPTG ($\mu M$) | Lux ($\mu M$) | Cin ($\mu M$) |
|---|---|---|---|---|---|
| | Condition A | 3.75 | 5 | 0.001 | 0.1 |
| | Condition B | 3.75 | 50 | 0.003 | 0.25 |
| | Condition C | 4.375 | 25 | 0.001 | 0.1 |
| | Condition D | 4.375 | 25 | 0.0025 | 0.7 |
| | Condition E | 6.5 | 25 | 0.001 | 0.1 |
| | Condition F | 5.5 | 50 | 0.001 | 0.1 |
| | Condition G | 6.5 | 25 | 0.0025 | 1 |
| | Condition H | 5.5 | 100 | 0.0025 | 0.25 |
| | Condition I | 9.5 | 25 | 0.002 | 0.1 |
| | Condition J | 10.5 | 25 | 0.002 | 0.1 |
| Condition K (Fig 6d) | | 18 | 500 | 0.008 | 12 |
| Condition L (Fig 6d) | | 18 | 500 | 0.008 | 4 |

Table S1: Conditions used for the 3-state composition control experiments. Molar concentrations of Salicylate (Bxb1), IPTG (Bxb1 RDF), Lux AHL (TP901) and Cin AHL (TP901 RDF) inducers employed for the experiments displayed in Figure 6c. The different color indexes corresponds to the different color traces in Figure S26.

.

### 1. Modeling 2-state phase variation in a chemostat

Let us first consider how one may model a single strain ($A$) growing through consumption of a substrate ($S$) in a chemostat environment. In abstracted terms one may expect the dynamics in Equation 1 for said species:

$$\frac{dA}{dt} = Growth - Death - Outflow$$

$$\frac{dS}{dt} = Substrate\ Inflow - Substrate\ Outflow + Substrate\ Consumption \tag{1}$$

$A$ experiences changes due to cell growth/division, death or outflow from the chemostat mechanism. Previous papers have shown individual cell death to be a rare event and to be the result of accumulated damage [1]. Because cells are consistently exiting the chemostat, we will assume death to be a negligible source of change in $A$. The levels of $S$ will change due to fresh media inflow, partially spent media outflow and consumption of substrate by cells for the process of replication. Due to the mechanism of a chemostat, this inflow term should be equal to the outflow term. We will now ascribe mathematical forms to each of these sources of change in Equation 2:

$$\frac{dA}{dt} = \left(\frac{\mu_a S}{K_a + S} - D\right) A$$

$$\frac{dS}{dt} = D(S_0 - S) - \left(\frac{\mu_a S}{K_a + S}\frac{A}{\gamma_a} + \frac{\mu_b S}{K_b + S}\frac{B}{\gamma_b}\right) \tag{2}$$

For the dynamics of $A$, cell growth is represented with a Monod model [2] and outflow is represented with a dilution term ($D$) applied to $A$. For the dynamics of $S$, substrate inflow and outflow are represented by the dilution term applied to the relevant substrate source (fresh, $S_0$ or spent media, $S$). Finally, substrate transformation into biomass is effectively the growth process from $\frac{dA}{dt}$ scaled by a yield term $\gamma_a$ which describes the efficiency by which substrate is converted to cell mass. We can now add a second strain, using the same exact dynamics as for the first strain, creating model that should exhibit competitive exclusion dynamics

$$\frac{dA}{dt} = \left(\frac{\mu_a S}{K_a + S} - D\right) A$$

$$\frac{dB}{dt} = \left(\frac{\mu_b S}{K_b + S} - D\right) B$$

$$\frac{dS}{dt} = D(S_0 - S) - \left(\frac{\mu_a S}{K_a + S}\frac{A}{\gamma_a} + \frac{\mu_b S}{K_b + S}\frac{B}{\gamma_b}\right)$$

We can introduce the phase variation dynamics associated with our 2-state circuit (highlighted in red). To model state transitions, we use simple linear switching terms ($\beta_j$) applied to the relevant species.

$$\frac{dA}{dt} = \left(\frac{\mu_a S}{K_a + S} - D\right) A + \beta_1 B - \beta_2 A$$

$$\frac{dB}{dt} = \left(\frac{\mu_b S}{K_b + S} - D\right) B + \beta_2 A - \beta_1 B$$

$$\frac{dS}{dt} = D(S_0 - S) - \left(\frac{\mu_a S}{K_a + S}\frac{A}{\gamma_a} + \frac{\mu_b S}{K_b + S}\frac{B}{\gamma_b}\right)$$

In this final model, $\mu_i$,$\gamma_i$ and $K_i$ are the maximum growth rate, yield constant and half-velocity constant for a given state $i$, respectively. $S_0$ and $D$, are the inlet substrate concentration and flow rate, respectively. Finally, $\beta_j$ is the switching rate associated with a given transition. Using non-dimensionalization we can reframe the system of equations and reduce the number of parameters to create the system in Equation 3:

$$\frac{d\mathbf{A}}{d\boldsymbol{t}} = \left( \frac{\boldsymbol{S}}{1+\boldsymbol{S}} - \overline{D} \right) \mathbf{A} + \overline{\beta_1}\mathbf{B} - \overline{\beta_2}\mathbf{A}$$

$$\frac{d\mathbf{B}}{d\boldsymbol{t}} = \left( \frac{\overline{\mu_b}\,\overline{K_b}\boldsymbol{S}}{1+\overline{K_b}\boldsymbol{S}} - \overline{D} \right) \mathbf{B} + \overline{\beta_2}\mathbf{A} - \overline{\beta_1}\mathbf{B}$$

$$\frac{d\boldsymbol{S}}{d\boldsymbol{t}} = \overline{D}\left(\overline{S_0} - \boldsymbol{S}\right) - \left( \frac{\boldsymbol{S}}{1+\boldsymbol{S}}\,\mathbf{A} + \frac{\overline{\mu_b}\,\overline{K_b}\boldsymbol{S}}{1+\overline{K_b}\boldsymbol{S}}\,\frac{\mathbf{B}}{\overline{\gamma_b}} \right) \tag{3}$$

$$\overline{D} = \frac{D}{\mu_a} \qquad \overline{\beta_j} = \frac{\beta_j}{\mu_a} \qquad \overline{\mu_b} = \frac{\mu_b}{\mu_a} \qquad \overline{K_b} = \frac{K_a}{K_b} \qquad \overline{S_0} = \frac{S_o}{K_a} \qquad \overline{\gamma_b} = \frac{\gamma_b}{\gamma_c}$$

For the simulations presented in Figure 2, the following non-dimensionalized parameters were used:

| Parameter | Uncontrolled Consortium | Phase-Varying Consortium |
|---|---|---|
| $\overline{\mu_b}$ | 1.3 | 1.3 |
| $\overline{K_b}$ | 1.0 | 1.0 |
| $\overline{\beta_1}$ | 0.0 | 0.2 |
| $\overline{\beta_2}$ | 0.0 | 0.1 |
| $\overline{D}$ | 0.5 | 0.5 |
| $\overline{S_0}$ | 9.0 | 9.0 |
| $\overline{\gamma_b}$ | 1.0 | 1.0 |

Table S2: 2-state phase variation parameter set. $\overline{K_b}$ and $\overline{\gamma_b}$ were set to unity as it is assumed both strains are of the same species and thus, bar growth rates, have similar growth properties. The values of $\overline{\beta_j}$ were selected in accordance with the expected timescales from a previous experimental study by Zhang et al [3]. $\overline{D}$ and $\overline{S_0}$ were selected to be in accordance with the expected functional regime of a chemostat system.

.

For the simulations used in Figure S1, the following parameters were used:

| Parameter | | | | | |
|---|---|---|---|---|---|
| $\overline{\beta_1}$ | 0.33 | 0.28 | 0.275 | 0.11 | 0.05 |
| $\overline{\beta_2}$ | 0.03 | 0.08 | 0.2 | 0.20 | 0.28 |

Table S3: 2-state composition tuning parameter set. Parameters used for the composition tuning simulations presented in Figure S1. The different color indexes corresponds to the different color traces in Figure S1. Unless otherwise specified, all other parameters used for these simulations can be found in Table S2

.

#### 1.1 Deriving Steady State Ratio of 2-state system

To simplify this derivation, we assume that both strains have the same yield constants and half-velocity constants i.e. $\overline{K_b} = \overline{\gamma_b} = 1$. At steady state, we can derive the following expressions:

$$0 = \left( X - \overline{D} - \overline{\beta_2} \right) r + \overline{\beta_1}$$

$$0 = \left( \overline{\mu_b} X - \overline{D} - \overline{\beta_1} \right) + \overline{\beta_2} r$$

$$X = \frac{\boldsymbol{S}}{1 + \boldsymbol{S}}, \quad r = \frac{\mathbf{A}^*}{\mathbf{B}^*}$$

We can solve for X for one expression and subsequently substitute it into the second expression, resulting in the following quadratic, with the solution in Equation 4:

$$r = \frac{\overline{D} + \overline{\beta_1} - \overline{\mu_b} \left( \overline{D} + \overline{\beta_2} - \frac{\overline{\beta_1}}{r} \right)}{\overline{\beta_2}}$$

$$0 = \overline{\beta_2} r^2 - \left[ (1 - \overline{\mu_b}) \overline{D} + \overline{\beta_1} - \overline{\mu_b}\, \overline{\beta_2} \right] r - \overline{\mu_b}\, \overline{\beta_1}$$

$$r = \frac{\alpha + \sqrt[2]{\alpha^2 + 4 \overline{\mu_b}\, \overline{\beta_1}\, \overline{\beta_2}}}{2 \overline{\beta_2}} \tag{4}$$

$$\alpha = (1 - \overline{\mu_b}) \overline{D} + \overline{\beta_1} - \overline{\mu_b}\, \overline{\beta_2}$$

The square root term will be strictly positive for all physical parameter values. Further this square root term will always be larger than $\alpha^2$ for positive $\overline{\mu_b}, \overline{\beta_1}, \overline{\beta_2}$ , hence for any system where switching occurs, we only need to consider the positive root. This implies the existence of single, unique positive $r$ i.e. coexistence for a given parameter set.

#### 1.2 Deriving the settling time of perturbations

We start by rewriting the system in matrix form in Equation 5:

$$\frac{d}{dt}\begin{bmatrix}\mathbf{A}\\ \mathbf{B}\end{bmatrix} = \begin{bmatrix} g_A & \overline{\beta_1} \\ \overline{\beta_2} & g_B \end{bmatrix}\begin{bmatrix}\mathbf{A}\\ \mathbf{B}\end{bmatrix} \tag{5}$$

$$g_a = \frac{\boldsymbol{S}}{1+\boldsymbol{S}} - \overline{D} - \overline{\beta_2},\ g_b = \frac{\overline{\mu_b}\boldsymbol{S}}{1+\boldsymbol{S}} - \overline{D} - \overline{\beta_1}$$

At steady-state we can derive a similar relationship to the previous section

$$0 = g_a\mathbf{A}^* + \overline{\beta_1}\mathbf{B}^*,\ 0 = \overline{\beta_2}\mathbf{A}^* + g_b\mathbf{B}^*$$

$$g_a = -\frac{\overline{\beta_1}}{r},\ g_b = -\overline{\beta_2}r$$

The characteristic polynomial of our Jacobian is the following and will have the eigenvalues displayed below.

$$\lambda^2 - (g_a + g_b)\lambda + \left(g_a g_b - \overline{\beta_1\beta_2}\right) = 0$$

$$\lambda_1 = 0,\ \lambda_2 = g_a + g_b = -\left(\frac{\overline{\beta_1}}{r} + \overline{\beta_2}r\right) < 0$$

The negative, real eigenvalue is relevant to the settling time of our system and implies deviations from steady state will decay exponentially (Equation 6).

$$\begin{aligned}\frac{\mathbf{A}}{\mathbf{B}}(t) - r &\sim exp(\lambda_2 t)\\ \tau_{settle} &= \frac{1}{|\lambda_2|} = \frac{1}{\left(\frac{\overline{\beta_1}}{r} + \overline{\beta_2}r\right)}\end{aligned} \tag{6}$$

When $r \approx 1$, both switching rates equally define the settling time. However, cases with bias towards one state ($r \neq 1$) will see the fast converting side of the pair weigh more heavily. For instance, if $r << 1$ then $\tau_{settle} \approx r/\overline{\beta_1}$, implying the speed of settling is largely determined by $\overline{\beta_1}$.

#### 2. Modeling 3-state phase variation in a chemostat

In this model, we employ the same basic chemostat model from the 2-state system and use the state transition dynamics we would expect to occur in the 3-state system (Fig. 5b, highlighted in red). We can model the dynamics of the respective genetics states ($A$, $B$, $C_A$ and $C_B$) and substrate ($S$) as follows:

$$\frac{dC_b}{dt} = \left( \frac{\mu_c S}{K_c + S} - D \right) C_b + \beta_4 B + \beta_2 C_a - C_b(\beta_1 + \beta_3)$$

$$\frac{dC_a}{dt} = \left( \frac{\mu_c S}{K_c + S} - D \right) C_a + \beta_4 A + \beta_1 C_b - C_a(\beta_3 + \beta_2)$$

$$\frac{dA}{dt} = \left( \frac{\mu_a S}{K_a + S} - D \right) A + \beta_1 B + \beta_3 C_a - A(\beta_2 + \beta_4)$$

$$\frac{dB}{dt} = \left( \frac{\mu_b S}{K_b + S} - D \right) B + \beta_3 C_b + \beta_2 A - B(\beta_1 + \beta_4)$$

$$\frac{dS}{dt} = D(S_0 - S) - \left( \frac{\mu_c S}{K_c + S} \frac{C_b}{\gamma_c} + \frac{\mu_c S}{K_c + S} \frac{C_a}{\gamma_c} + \frac{\mu_a S}{K_a + S} \frac{A}{\gamma_a} + \frac{\mu_b S}{K_b + S} \frac{B}{\gamma_b} \right)$$

$\mu_i$,$\gamma_i$ and $K_i$ are the maximum growth rate, yield constant and half-velocity constant for a given state $i$, respectively. $S_0$ and $D$, are the inlet substrate concentration and flow rate, respectively. Finally, $\beta_j$ is the switching rate associated with a given transition. Note that states $C_A$ and $C_B$ adopt the same protein state/phenotype, and they thus have identical growth parameters in spite of their distinct genetic states. Using non-dimensionalization we can reframe the system of equations and reduce the number of parameters as follows, yielding Equation 7:

$$\frac{d\mathbf{C}_b}{d\boldsymbol{t}} = \left( \frac{\boldsymbol{S}}{1+\boldsymbol{S}} - \overline{D} \right) \mathbf{C}_b + \overline{\beta_4}\mathbf{B} + \overline{\beta_2}\mathbf{C}_a - \mathbf{C}_b \left( \overline{\beta_1} + \overline{\beta_3} \right)$$

$$\frac{d\mathbf{C}_a}{d\boldsymbol{t}} = \left( \frac{\boldsymbol{S}}{1+\boldsymbol{S}} - \overline{D} \right) \mathbf{C}_a + \overline{\beta_4}\mathbf{A} + \overline{\beta_1}\mathbf{C}_b - \mathbf{C}_a \left( \overline{\beta_3} + \overline{\beta_2} \right)$$

$$\frac{d\mathbf{A}}{d\boldsymbol{t}} = \left( \frac{\overline{\mu_a}\,\overline{K_a}\boldsymbol{S}}{1+\overline{K_a}\boldsymbol{S}} - \overline{D} \right) \mathbf{A} + \overline{\beta_1}\mathbf{B} + \overline{\beta_3}\mathbf{C}_a - \mathbf{A} \left( \overline{\beta_2} + \overline{\beta_4} \right)$$

$$\frac{d\mathbf{B}}{d\boldsymbol{t}} = \left( \frac{\overline{\mu_b}\,\overline{K_b}\boldsymbol{S}}{1+\overline{K_b}\boldsymbol{S}} - \overline{D} \right) \mathbf{B} + \overline{\beta_3}\mathbf{C}_b + \overline{\beta_2}\mathbf{A} - \mathbf{B} \left( \overline{\beta_1} + \overline{\beta_4} \right)$$

$$\frac{d\boldsymbol{S}}{d\boldsymbol{t}} = \overline{D}\left( \overline{S_0} - \boldsymbol{S} \right) - \left( \frac{\boldsymbol{S}}{1+\boldsymbol{S}}\mathbf{C}_b + \frac{\boldsymbol{S}}{1+\boldsymbol{S}}\mathbf{C}_a + \frac{\overline{\mu_a}\,\overline{K_a}\boldsymbol{S}}{1+\overline{K_a}\boldsymbol{S}}\frac{\mathbf{A}}{\overline{\gamma_a}} + \frac{\overline{\mu_b}\,\overline{K_b}\boldsymbol{S}}{1+\overline{K_b}\boldsymbol{S}}\frac{\mathbf{B}}{\overline{\gamma_b}} \right) \quad (7)$$

$$\overline{D} = \frac{D}{\mu_c} \qquad \overline{\beta_j} = \frac{\beta_i}{\mu_c} \qquad \overline{\mu_i} = \frac{\mu_i}{\mu_c} \qquad \overline{K_i} = \frac{K_c}{K_i} \qquad \overline{S_0} = \frac{S_o}{K_c} \qquad \overline{\gamma_i} = \frac{\gamma_i}{\gamma_c}$$

We can numerically integrate this system of differential equations to investigate the properties of our proposed Timesharing design in a chemostat environment. Further, to simulate the conventional, uncontrolled case, all switching parameters ($\beta_i$) can be set to zero.

For the set of simulations in Fig. 5c, the following parameters were used:

| Parameter | Phase-Varying Consortium |
|---|---|
| $\overline{\mu_a}$ | 1.1 |
| $\overline{\mu_b}$ | 1.3 |
| $\overline{K_a}$ | 1.0 |
| $\overline{K_b}$ | 1.0 |
| $\overline{\beta_1}$ | 0.145 |
| $\overline{\beta_2}$ | 0.090 |
| $\overline{\beta_3}$ | 0.1 |
| $\overline{\beta_4}$ | 0.062 |
| $\overline{D}$ | 0.5 |
| $\overline{S_0}$ | 10.0 |
| $\overline{\gamma_a}$ | 1.0 |
| $\overline{\gamma_b}$ | 1.0 |

Table S4: 3-state phase variation parameter set. Simulations were performed for 60 divisions. $\overline{K_i}$ and $\overline{\gamma_i}$ were set to unity as it assumed both strains are of the same species and thus, bar growth rates, have similar growth properties. The values of $\overline{\beta_j}$ were selected in accordance with the expected timescales from a previous experimental study by Zhang et al [3]. $\overline{D}$ and $\overline{S_0}$ were selected to be in accordance with the expected functional regime of a chemostat system.

.

#### 2.1 Parameter sweep for 3-state system

To probe the composition control property of our 3-state model, we performed a parameter sweep across the $\beta j$ values (Figure S12). For these simulations we separated our switching rates into two paired sets, $\beta_1, \beta_2$ and $\beta_3, \beta_4$. We constrained that each paired set must sum to a value of 0.4 and varied the values of a given switching rate within a pair from 0 to 0.4 (40 intervals), creating $40^2$ unique parameter sets. We then simulated our system for 45 divisions with each of these parameter sets, letting it reach steady state and plotting this steady state composition on our ternary diagram.

### 3. Simulations of 2-state mother machine experiments

During the mother machine experiments, it was noted that the observed YFP and mTurquoise residence times had distinct distribution shapes (Fig. 3d). Notably, the YFP residence times has a very distinct shape with an evenly distributed probability density across the function. We posited that due to the limited observation time associated with our mother machine experiments (22 hours), we were never able to observe any residence time longer than 22 hours. Effectively, we were sampling a truncated (at 22 hours) distribution of residence times, resulting in an incomplete and biased distribution. We developed a model of our system in the mother machine to explore how varying observation time of a switching system may influence the observed distribution of residence times.

#### 3.1 Model Framework

Our simulations rely on two objects implemented in python. Below is a brief description of the most important methods and attributes associated with each object:

- `Cell`: This object represents a single cell in the mother machine. It has `state` and `state counter` attributes that allows the `Cell` object to exist in either a `yellow` or `blue` state and track how long it has existed in that state. A `Cell` object can transition states using the `transition` method and record switching events in the `arrival time` attribute. Transitions are controlled via a Bernoulli process, with switching rates input being scaled to the timestep size to generate switching probabilities for the Bernoulli process. Finally, a `label` attribute define cells as either `mothers` or `daughters` and daughters cells can be generated via a `spawn` method

- `Cell_env`: This object acts a simulation environment in which many `Cell` objects can be run over many timesteps. It contains all `Cell` objects in a `running_list` attribute and tracks the dynamics in a `history` pandas dataframe attribute. It has methods to run multiple time-dependent process asynchronously, including transitions (`run_trans`) and spawning (`run_spawn`). Critically, the maximum number of cells is capped via a `max_pop` attribute. If the number of `Cell` objects exceeds this cap, the `cull` method prunes `running_list` to the cap, retaining objects with the `mother` label over objects with the `daughter` label.

The code required to run simulations and an accompanying notebook with the relevant simulations and analysis can be found in the associated GitHub repository.

#### 3.2 Model Results

We modeled the yellow trace from Fig. 4b (0 uM Sal, 250 uM IPTG) and the associated residence time distribution from Fig. 3d. We assumed the mTurquoise residence times to be accurate and then determined the expected YFP residence times based off the steady state composition of the experiment. This led to the following parameter set for the experiment in question.

| Parameter Name | Value |
|---|---|
| `max_pop` | 5000 cells |
| `to_blue_probablity` | 0.0057 |
| `to_yellow_probability` | 0.057 |

Table S5: Parameters used for mother machine simulation. Probabilities were determined by fixing the simulation timescale to 10 minutes, the acquisition interval of our experiments. In this context, the average switches (to a given state) per 10 min was used as these relevant state transition probabilities in a Bernoulli process

.

We then simulated this parameter set across four sets of observation times (22, 48, 192, 768 hours), plotting the YFP and mTurquoise residence times separately.

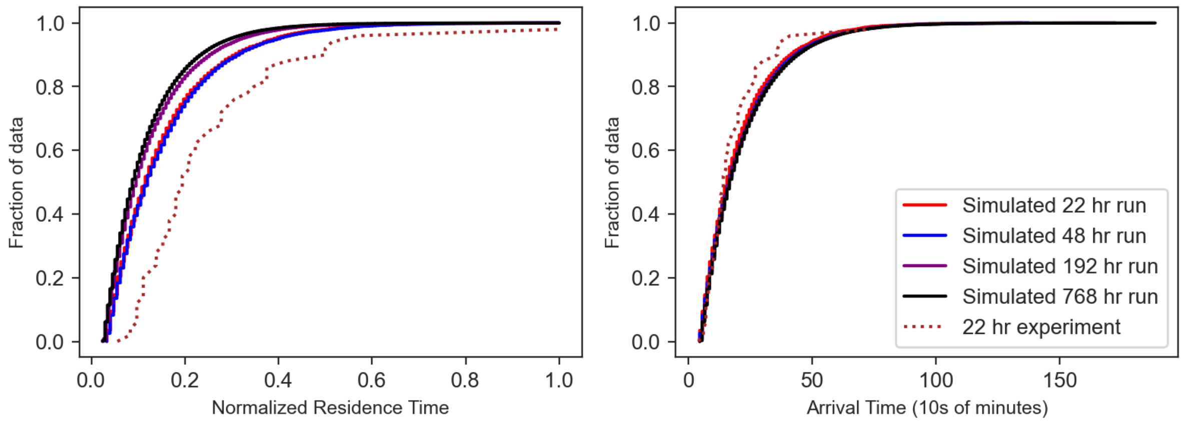


Figure S27: Simulated and experimental mTurquoise residence times. a. Normalized residence time distributions. Max observed residence time of a given sample is used to normalize the residence times, in practice this is how an experimentally collected cumulative distribution function would look. b. Raw residence time distributions. No normalization of residence times.

We first examine the mTurquoise residence times. Due to the fast switching rates associated with the mTurquoise to YFP transition, observation time does not have a drastic impact on the shape of the residence time distribution. The shape of these simulated distributions is quite similar to the shape of experimentally observed distribution.

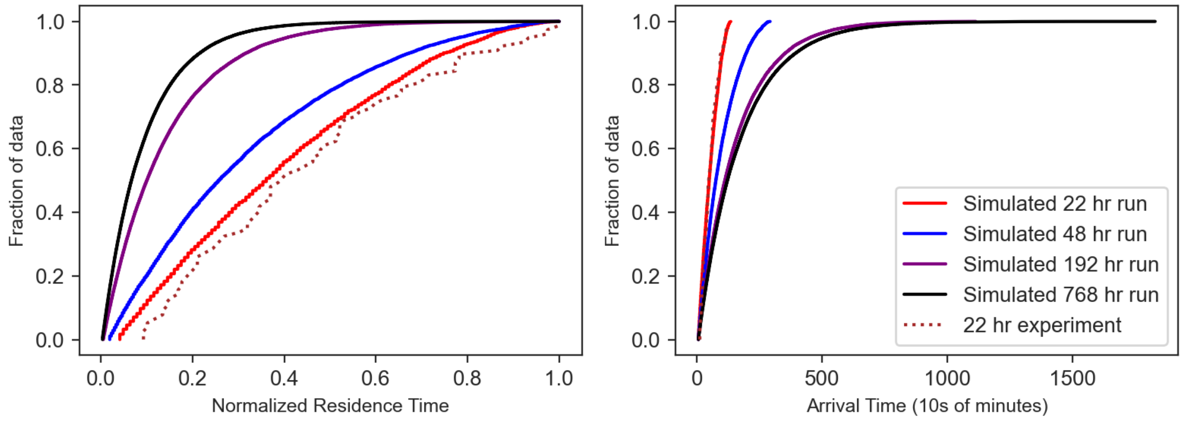


Figure S28: Simulated and experimental YFP residence times. a. Normalized residence time distributions. Max observed residence time of a given sample is used to normalize the residence times, in practice this is how an experimentally collected cumulative distribution function would look. b. Raw residence time distributions. No normalization of residence times.

We now examine the YFP residence times. We expect that the much slower switching rates associated with the YFP to mTurquoise transition will lead to many residence times that occur past the 22 hr time limit of our experiment. As we can see, the shape of the residence time distribution drastically shifts as the observation time increases. Further, we note that short observations samples e.g. 22 hr and 48 hr have shapes very similar to the experimentally observed distribution. These simulations support the hypothesis that observation time must be extended to fully capture the residence time distribution, however experiments recreating these simulation results would be necessary to fully confirm this hypothesis.

### References

1. Wang, P. *et al.* Robust Growth of Escherichia coli. English. *Current Biology* 20. Publisher: Elsevier, 1099–1103. ISSN: 0960-9822. https://www.cell.com/current-biology/abstract/S0960-9822(10)00524-5 (2025) (June 2010).

2. Monod, J. *Recherches sur la croissance des cultures bactériennes* (Hermann & cie, Paris, 1942).

3. Zhang, Q., Azarin, S. M. & Sarkar, C. A. Model-guided engineering of DNA sequences with predictable site-specific recombination rates. en. *Nat Commun* 13. Publisher: Nature Publishing Group, 4152. ISSN: 2041-1723. https://www.nature.com/articles/s41467-022-31538-3 (2025) (July 2022).
